# Allosteric effector ppGpp potentiates the inhibition of transcript initiation by DksA

**DOI:** 10.1101/188680

**Authors:** Vadim Molodtsov, Elena Sineva, Lu Zhang, Xuhui Huang, Michael Cashel, Sarah E. Ades, Katsuhiko S. Murakami

**Affiliations:** Department of Biochemistry and Molecular Biology, The Center for RNA Molecular Biology, The Pennsylvania State University, University Park, PA 16802, USA; State Key Laboratory of Structural Chemistry, Fujian Institute of Research on the Structure of Matter, Chinese Academy of Sciences, Fuzhou, Fujian, 350002, China; Department of Chemistry, The Hong Kong University of Science and Technology, Clear Water Bay, Kowloon, Hong Kong; Intramural Research Program, Eunice Kennedy Shriver, National Institute of Child Health and Human Development, National Institutes of Health, Bethesda, MD 20892, USA

## Abstract

DksA and ppGpp are the central players in the *Escherichia coli* stringent response and mediate a complete reprogramming of the transcriptome from one optimized for rapid growth to one adapted for survival during nutrient limitation. A major component of the response is a reduction in ribosome synthesis, which is accomplished by the synergistic action of DksA and ppGpp bound to RNA polymerase (RNAP) inhibiting transcription of rRNAs. Here, we report the X-ray crystal structures of *E. coli* RNAP holoenzyme in complex with DksA alone and with ppGpp. The structures show that DksA accesses the template strand at the active site and the downstream DNA binding site of RNAP simultaneously and reveal that binding of the allosteric effector ppGpp reshapes the RNAP–DksA complex. The structural data support a model for transcriptional inhibition in which ppGpp potentiates the destabilization of open complexes on rRNA promoters by DksA. We also determined the structure of RNAP–TraR complex, which reveals the mechanism of ppGpp-independent transcription inhibition by TraR. This work establishes new ground for understanding the pleiotropic effects of DksA and ppGpp on transcriptional regulation in proteobacteria.

**Highlights:**

- DksA has two modes of binding to RNA polymerase
- DksA is capable of inhibiting the catalysis and influences the DNA binding of RNAP
- ppGpp acts as an allosteric effector of DksA function
- ppGpp stabilizes DksA in a more functionally important binding mode

## INTRODUCTION

The stringent response in bacteria is a global rearrangement of cellular metabolism from one optimized for vegetative growth to one optimized for stress survival and is accompanied by changes in the expression of over 500 genes including those involved in ribosome biogenesis, amino acid synthesis, virulence, survival during host invasion, antibiotic resistance and persistence (Dalebroux and Swanson, 2012; Durfee et al., 2008; Hauryliuk et al., 2015; Potrykus and Cashel, 2008). Accumulation of unusual nucleotides, the bacterial alarmones ppGpp (guanosine tetraphosphate) and pppGpp (guanosine pentaphosphate), here referred to collectively as ppGpp, in response to nutrient downshifts triggers the stringent response. In the proteobacteria, ppGpp exerts its effects on gene expression primarily by regulating the activity of RNA polymerase (RNAP) during transcription initiation.

Transcriptional regulation by ppGpp has been studied most extensively in *E. coli.* Unlike DNA-binding transcriptional regulators, which exert their effects only on genes with regulator binding sites properly positioned relative to the promoter, ppGpp binds to RNAP and has the potential to influence the expression of any gene. Whether expression of a given gene is altered by ppGpp depends on the properties of the promoter directing expression of that gene (Haugen et al., 2008). Many genes are unaffected by ppGpp while others are repressed, notably those encoding stable rRNAs, or activated, such as those encoding genes for amino acid biosynthesis.

ppGpp often works in conjunction with DksA, a 17.5 kDa protein conserved in proteobacteria. DksA belongs to the class of RNAP secondary channel binding transcription factors that includes GreA and GreB (Opalka et al., 2003; Sekine et al., 2015), Gfh1 (Tagami et al., 2010) and TraR (Blankschien et al., 2009b) (**Figs. S1A, B and C**). The crystal structure of DksA showed that it consists of five α helixes organized into three structural parts (Perederina et al., 2004). The globular domain (G domain) is formed by amino acids from residues 1-32 and 109-134 and includes two α helixes (α1 and α4) along with the Zn binding region. The central segment of the polypeptide chain (residues 33-108) forms an extended coiled-coil domain (CC domain) with two long α-helices (α2 and α3) connected by a short linker (CC tip). The C-terminal end of DksA forms an extended α helix (CT-helix, residues 135-151), which is loosely connected to the rest of the protein. The structural organization of DksA is similar to that of the Gre and Gfh1, which also contain the G and CC domains, however, the CT-helix is present only in DksA.

The cellular concentration of DksA remains constant under different growth conditions, consequently ppGpp serves as the signal of nutrient limitation (Paul et al., 2004). The importance of DksA for ppGpp activity is demonstrated by the finding that regulation of rRNA and amino acid promoter expression in response to growth rate and nutrient limitation are lost in a Δ*dksA* strain, similar to a ppGpp null strain (Paul et al., 2004; Paul et al., 2005). In vitro, DksA and ppGpp act synergistically to regulate transcription at both stable RNA promoters and amino acid biosynthetic promoters, consistent with the in vivo observations (Paul et al., 2004; Paul et al., 2005). Together, they decrease the life time of open complexes (RPo) formed by RNAP on all promoters, but the outcome of this destabilization depends on the promoter. For those promoters that form intrinsically short-lived RPo, such as the stable RNA promoters, further reduction in stability favors accumulation of the closed complex (RPc) and dissociation of RNAP from the promoter, ultimately resulting in inhibition of transcription. Although DksA and ppGpp also reduce the stability of promoter forming long-lived RPo, it is not limiting for transcription initiation and these promoters are not inhibited. Therefore, it appears that a defining characteristic of promoter subject to negative regulation by ppGpp and DksA is formation of an intrinsically unstable RPo. The mechanism of positive regulation by ppGpp and DksA is less well understood and several mechanisms have been suggested. For the *hisG* promoter, these factors were found to increase the isomerization rate from RPc to RPo (Paul et al., 2005), while at the *uspA* promoter ppGpp and DksA increased the rate of promoter clearance by destabilizing the intrinsically stable RPo (Gummesson et al., 2013).

Two binding sites for ppGpp have been identified on *E. coli* RNAP. Site 1 lies in a cavity formed by the β’ and ω subunits (Mechold et al., 2013; Ross et al., 2013; Zuo et al., 2013) and ppGpp binds independently of DksA at this site. RNAP reconstituted without the ω subunit is no longer inhibited by ppGpp in vitro in the absence of DksA (Igarashi et al., 1989; Vrentas et al., 2005). However, in the presence of DksA, sensitivity of the Δω RNAP to ppGpp is restored (Vrentas et al., 2005). Deletion of the *E. coli* gene encoding ω *(rpoZ)* resulted in relatively mild phenotypes in vivo (Gentry et al., 1991), suggesting that site 1 alone is not sufficient for induction of the stringent response. A recent study identified another ppGpp binding site near the secondary channel of RNAP, which is ~60 Å away from site 1 (Ross et al., 2016). This second ppGpp binding site (site 2) is formed only in the presence of DksA. The growth rate of an *E. coli* strain with a variant of RNAP carrying mutation that disrupts site 2 is severely impaired in minimal medium as would be expected if ppGpp binding at site 2 were primarily responsible for reprogramming gene expression during the stringent response (Ross et al., 2016).

The TraR transcription regulator binds in the secondary channel of RNAP similar to DksA and can regulate transcription in a manner analogous to DksA (Blankschien et al., 2009b; Gopalkrishnan et al., 2017; Grace et al., 2015). TraR is not a chromosomal gene and is primarily carried on conjugative plasmids that enable horizontal gene transfer. The full spectrum of TraR functions in cell metabolism remains to be determined, but it has been proposed to play a role in bacterial antibiotic resistance, pathogenicity and virulence (Maneewannakul and Ippen-Ihler, 1993). TraR is a truncated version of DksA lacking the first 68 amino acids. Despite its shorter length, TraR can inhibit transcription of rRNA genes and activate transcription of amino acid biosynthesis genes (Gopalkrishnan et al., 2017). Interestingly, TraR function is not influenced by ppGpp and its transcriptional regulatory activities are more closely resemble those of DksA with ppGpp, than DksA alone.

In this study, we report crystal structures of the *E. coli* RNAP σ^70^ holoenzyme—DksA complex with and without ppGpp. The structures reveal the precise interaction network among RNAP, DksA and ppGpp, and provide structural basis for destabilization of RPo by these factors. Conformational changes are observed in both RNAP and DksA in the binary complex (RNAP—DksA) and the ternary complex (RNAP—DksA/ppGpp) that have implications for the mechanism of transcriptional regulation. Importantly, the location of DksA is altered by ppGpp, demonstrating that ppGpp allosterically potentiates the function of DksA. We also determined the crystal structure of RNAP in complex with TraR and established the mechanism by which this small secondary channel binding protein effectively regulates transcription without ppGpp.

## RESULTS

### Structure of the RNAP—DksA binary complex

Because the secondary channel of the *E. coli* RNAP σ^70^ holoenzyme is widely open and accessible in the crystalline state (Murakami, 2013), we were able to prepare co-crystals of the RNAP—DksA complex by soaking DksA into preformed RNAP crystals (**Table 1**). A similar approach was used for reconstitution and X-ray crystal structure determination of yeast RNAP II with the transcription factor TFIIS, which also binds at the secondary channel (Kettenberger et al., 2003). It is interesting to note that the crystal lattice can tolerated large structural rearrangements induced by DksA, and co-crystal structures were obtained from crystals with the same form and packing as RNAP alone. As such, the detected structural changes can be attributed to DksA binding to RNAP and not to changes in crystal packing. We also determined the crystal structures of RNAP—DksA/ppGpp and RNAP–TraR complexes by soaking these factors into preformed RNAP crystals (**see below**). Although the binding sites of DksA, ppGpp and TraR are accessible in the *E. coli* RNAP σ^70^ holoenzyme crystal and certain conformational changes of RNAP and DksA are observed in this study, it is possible that the full range of conformational changes are not observed here due to crystal lattice constraint.

Crystal structures of RNAP (Murakami, 2013) and DksA (Perederina et al., 2004) were fitted to the electron density map without ambiguity resulting in a structure at 4.5 Å resolution (**Fig. 1 and SFig. S2A**). As predicted from biochemical and modeling experiments (Lennon et al., 2012; Parshin et al., 2015), DksA is located in the secondary channel of RNAP. The electron density map of DksA was well defined in the complex with the exception of G domain where the density was scattered and discontinuous (**SFig. S1D**), indicating that this part of DksA is not stably bound to RNAP. However, we were able to dock the DksA structure into the density that was obtained and model the likely position of the G domain with RNAP. DksA primarily contacts the β’ subunit and all three domains (CC, G, and CT-helix) are involved in the interaction with RNAP (**Fig. 1, Movie S1**).

**Figure 1.**
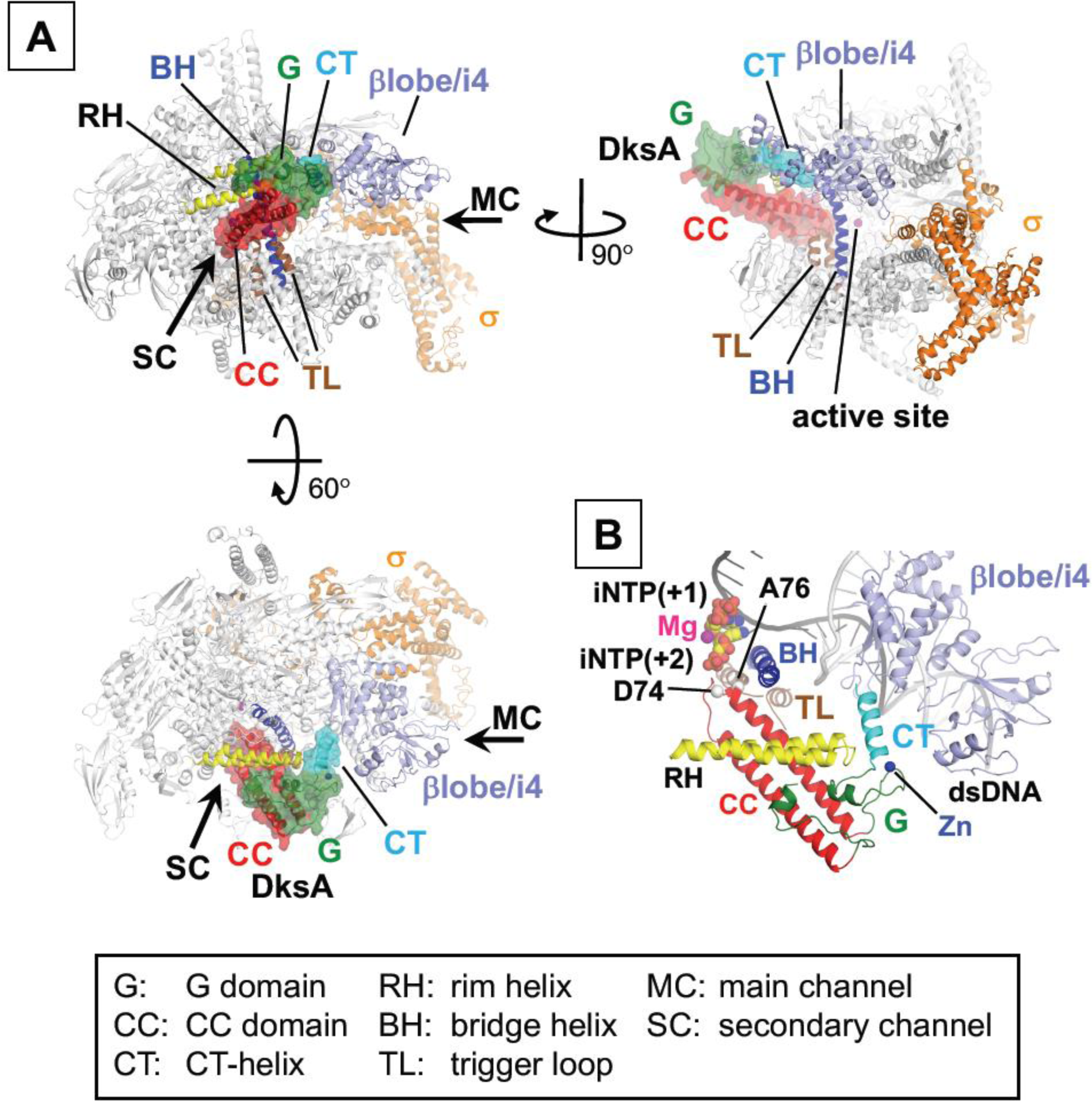
Structure of RNAP—DksA binary complex. (**A**) RNAP is depicted as ribbons and DksA is depicted as a ribbon with partially transparent surface. Relative orientations of proteins in the panels are indicated. Domains, motifs and regions of RNAP and DksA discussed in the text are labeled. (**B**) A model of DksA bound to RPo. RNAP and DksA are depicted as ribbons, and positions of D74 and A76 residues of DksA are indicated. The DNA and iNTPs bound at +1 and +2 positions are modeled using *T. thermophilus* transcription initiation complex (PDB: 4Q4Z) (Basu et al., 2014). Orientation of RNAP in this panel is the same as in the left-bottom panel in **A**.

The DksA CC domain is positioned in a cleft on RNAP that runs from the secondary to the main channels. One surface of the DksA CC domain faces the β’rim helix and runs alongside it but makes few physical interactions, allowing the CC domain to move its position after the ppGpp binding at site 2 (**see below**). The opposite surface of CC domain faces β’i6 (also known as β’SI3) (Chlenov et al., 2005), a lineage-specific insertion in the middle of trigger loop. β’i6 was not resolved in any of the structures determined in this study indicating that it remains flexible. This observation is consistent with the proposal that DksA only binds to forms of RNAP in which β’i6 is mobile (Furman et al., 2013). The CC tip, which is required for DksA function (Lee et al., 2012), inserts into the secondary channel and comes within ~16 Å of the catalytic Mg^2+^ coordinated at the active site (**Fig. 1B**). Amino acid substitutions at the CC tip (D74 and A76) eliminate DksA function without affecting its apparent affinity for RNAP (Lee et al., 2012; Parshin et al., 2015). These residues are near the bridge helix (BH), but do not directly contact the BH. Finally, the CC domain is positioned in a manner that would cause a steric class when trigger loop folds (**Fig. S3A**), presenting a problem for the catalytic step in RNA synthesis which requires the trigger loop folding (Yuzenkova et al., 2010; Zhang et al., 2010). Interestingly, the CC tip can be crosslinked to the trigger loop (Lennon et al., 2012) providing further evidence that an interplay between these elements may be involved in regulation of transcription by DksA.

In addition to the CC domain, the CT-helix and G domain make contacts with RNAP. The CT-helix extends toward the βlobe/i4 domain (also known as βSI1), which forms one of pincers surrounding the main DNA binding channel (**Fig. 1B**). Direct contact with the CT-helix rotates the βlobe/i4 domain as a rigid body toward the β’rim helix (**Fig. 2C, Movie S2**). Deletion of the CT-helix or of the βlobe/i4 domain disrupt DksA binding to RNAP and transcriptional repression underscoring the importance of these interactions (Parshin et al., 2015). Additional contacts are formed between the G domain and the edge of β’rim helix resulting in a 7.2 Å shift in the rim helix position compared with the apo-form RNAP. The conformation of DksA itself also changes upon RNAP binding; the α1, α4 and CT-helix move in different directions from the center of G domain, weakening the hydrophobic packing between the CC and G domains (**Fig. 2F, Movie S3**).

**Figure 2.**
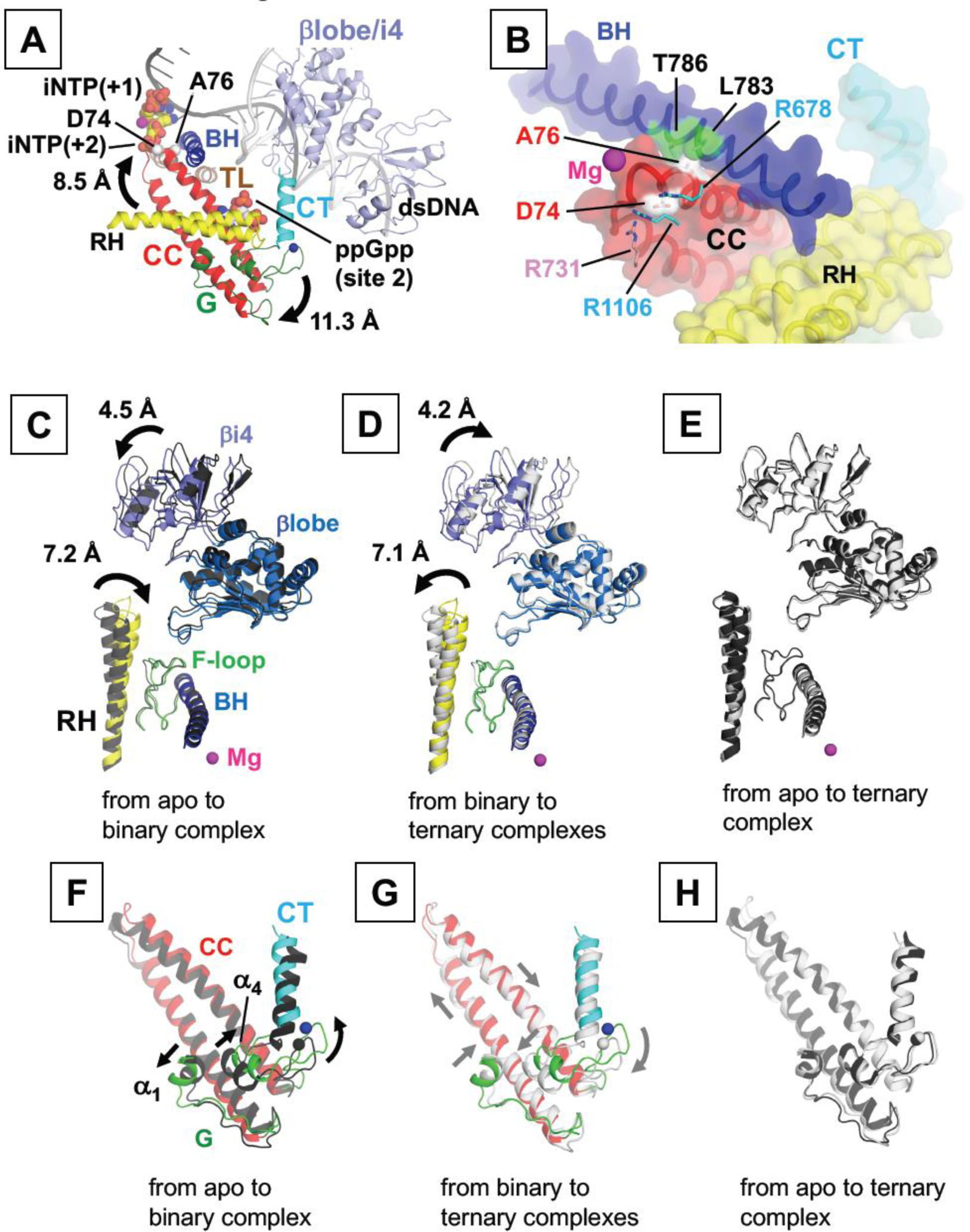
Conformation changes in DksA and RNAP induced by DksA binding alone and with ppGpp. (**A**) A model of DksA and ppGpp bound to RPo. RNAP, DksA, DNA and iNTPs are depicted as in Fig. 1B. ppGpp bound at the site 2 is shown in CPK representation. Rotation of DksA induced by ppGpp binding is indicated by the arrows. (**B**) A magnified view of the CC tip of DksA in the ternary complex. DksA, BH and RH of RNAP are depicted as ribbons with partially transparent surfaces. The D74 and A76 residues of DksA and amino acid residues of RNAP contacting with these residues are depicted as sticks and labeled. (**C-E**) Conformational changes in RNAP upon binding of DksA (**C**) and ppGpp (**D**). Movement of the β’rim helix and βlobe/i4 domain is indicated by arrows. RNAP in the binary complex is shown in color, while RNAPs in the apo-form and in the ternary complex are shown as black and white ribbons, respectively. (**E**) Superimposed ribbon representation of RNAP between its apo-form (black) and in the ternary complex (white). (**F-H**) Conformational changes in DksA upon binding to RNAP alone (**F**) and with ppGpp (**G**). Movements of DksA are indicated by arrows. DksA in the binary complex is shown in the same color as in **A**, and DksA molecules in the apo-form and in the ternary complex are shown as black and white ribbons, respectively. (**H**) Superimposed ribbon representation of DksA in the apo-form (black) and in the ternary complex (white).

### Structure of the RNAP—DksA/ppGpp ternary complex

We prepared co-crystals of the ternary complex containing RNAP, DksA and ppGpp in the same manner as the RNAP—DksA complex, by stepwise soaking of DksA and then ppGpp into preformed RNAP crystals. As with the DksA-RNAP co-crystals, the crystal form and packing remained the same and we were able to determine the structure to 4.3 Å resolution (**Table 1, Movie S4**). Electron densities corresponding to ppGpp molecules were found at the locations previously described for ppGpp binding site 1 (Mechold et al., 2013; Ross et al., 2013; Zuo et al., 2013) and site 2 (Ross et al., 2016) (**Fig. S2B, see below**).

ppGpp binding sites 1 and 2 are separated from each other by 62 Å, and by 31 Å and 40 Åfrom the active site, respectively (**Fig. 3A**) indicating that they do not function via direct interactions with each other or the active site. The chemical environment of ppGpp binding site 1 is distinct from that of site 2. At site 1, ppGpp binds in a shallow pocket at the interface between the β’ and ω subunits. One surface of ppGpp binds to RNAP while the other is exposed to solvent (**Fig. 3B**). Although the orientation of individual side chains around the ppGpp binding sites could not be resolved at the resolution of the current structure, several amino acid residues are located in positions where they are likely to directly contact ppGpp. The guanine base is sandwiched by side chains of R362 and I619 of the β’ subunit in addition to facing H364 and D622 residues, likely forming with them a hydrogen bond and salt bridge, respectively.

**Figure 3.**
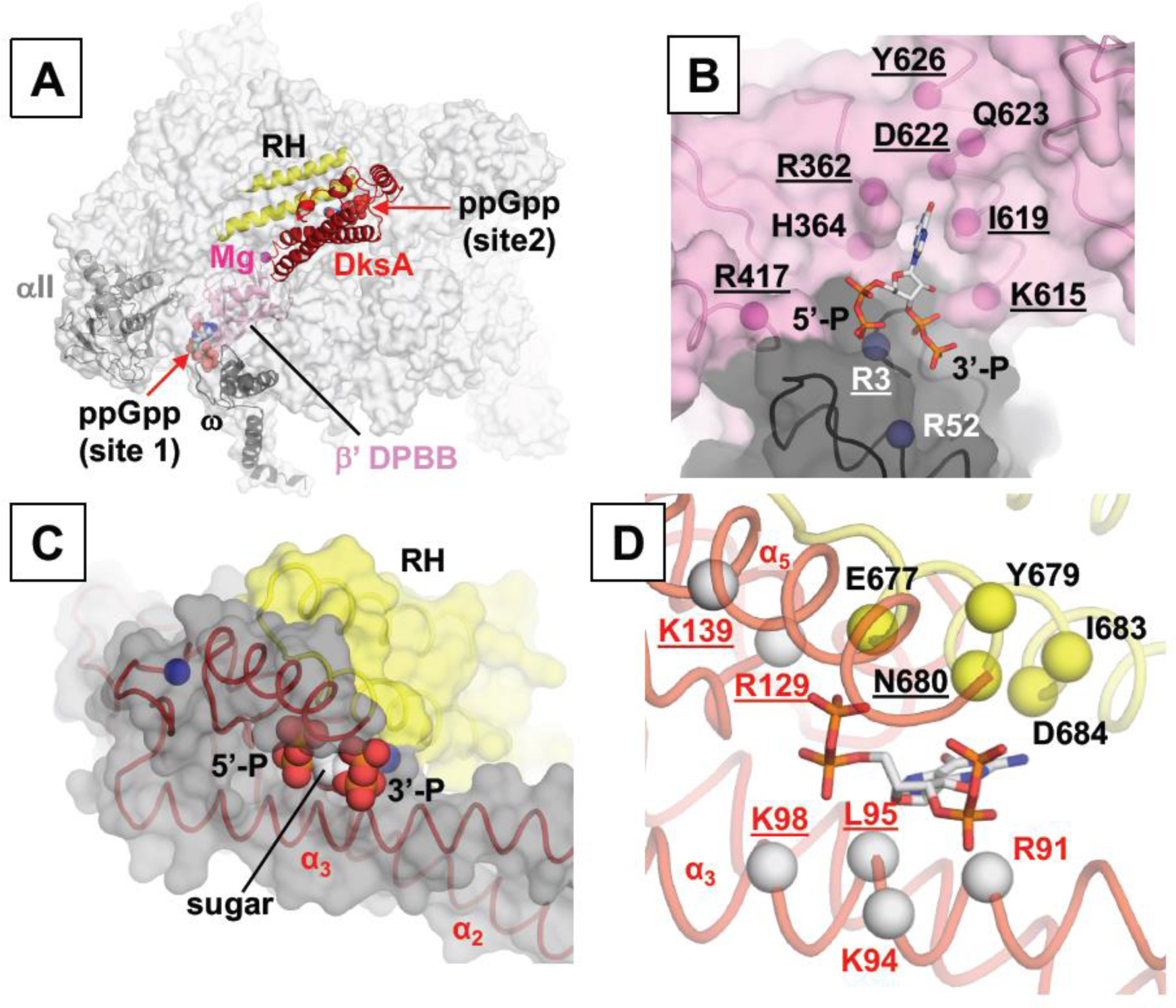
ppGpp binding sites on RNAP. (**A**) ppGpp binding sites 1 and 2 are shown in the RNAP—DksA/ppGpp complex (RNAP: transparent white surface; DksA: red ribbon; ppGpp: CPK surface). Domains and subunits of RNAP forming the ppGpp binding sites are shown as ribbon models and labeled. (**B**) ppGpp binding site 1 is shown. The β’ (pink) and ω (dark gray) subunits are depicted as ribbons with transparent surfaces and ppGpp as a stick representation. Positions of amino acid residues involved in ppGpp binding are indicated as spheres and labeled. Residues determined from biochemical and genetic studies to be important for ppGpp binding are underlined (Ross et al., 2013). (**C**) ppGpp binding site 2 is shown. DksA (gray surface and red ribbon), β’rim helix (yellow surface) and ppGpp (CPK surface) are shown. Positions of the ribose sugar and 5’-and 3’-phosphate groups of ppGpp are indicated. (**D**) Positions of amino acid residues involved in ppGpp binding are indicated as spheres (β’: yellow, DksA: red) and labeled. Residues determined from biochemical and genetic studies (Parshin et al., 2015; Ross et al., 2016) to be important for ppGpp binding are underlined. The orientations of panels **C** and **D** are the same.

ppGpp binding site 2 is formed by a narrow cleft at the interface between DksA and the β’rim helix. The guanine base of ppGpp inserts into the cleft, while the sugar and phosphate groups remain solvent exposed (**Fig. 3C**). DksA primarily interacts with phosphate groups of ppGpp. Basic residues found in the CC domain (R91, K94, K98) and the CT-helix (K139) are in close proximity to the 3’ or 5’ phosphates and likely form salt bridges with ppGpp (**Fig. 3D**). The L95 side chain is positioned such that it may interact with the guanine base via a van der Waals interaction. In the CT-helix, R129 is close to E677 of β’ subunit and could neutralize its charge, which is located near the 5’ diphosphate group of ppGpp. In addition to the contacts between DksA and the phosphates, the β’ rim helix interacts with the guanine base via a salt bridge (D684), a hydrogen bond (N680) and van der Waals contacts (Y679, N680 and I683). Biochemical assays support the structural findings. DksA L95 K98, R129 and K139 have been shown to be critical for ppGpp binding and activity, but do not affect DksA activity. Mutation of DksA R91 also strongly reduces ppGpp binding. In RNAP, mutation of β’ D684 and N680 significantly reduced ppGpp binding at site 2 without altering binding of and regulation by DksA alone, while mutation of β’ E677 to alanine eliminated binding and regulation of both DksA and ppGpp. Residues implicated as critical for DksA and ppGpp activity at site 2 from previous studies (Parshin et al., 2015; Ross et al., 2016) are underlined in **Fig. 3C**.

After the ppGpp binding, DksA undergoes a rigid body rotation centered around the ppGpp binding site (**Fig. 2A, Movie S5**) leading to deeper insertion of the CC tip into the secondary channel that positions the D71 side chain adjacent to the NTP binding site (*i*+1 site) of the RNAP active site. D71 may also form salt bridges with R678/R1106 (β subunit) and R731 (β’ subunit) residues (**Fig. 2B**). Mutation of R678 and R1106 to alanine has been shown to reduce regulation and destabilization of RPo by DksA (Parshin et al., 2015). The ppGpp-induced rotation of DksA also brings the CC tip into contact with the BH such that the A76 residue faces the center of the BH (amino acids 775-790) and fits snugly in a cavity surrounded by BH residues L783 and T786 (**Fig. 2B**). This direct contact between the BH and the CC domain of DksA may influence the stability of RPo (**see Discussion: The structural basis for transcription inhibition by DksA/ppGpp and TraR**).

Interestingly, the conformational changes in both RNAP and DksA triggered by DksA binding in the absence of ppGpp were not observed in the ternary structure. The β’rim helix and the βlobe/i4 domain were not distorted and assumed the same conformations seen in the apo form RNAP (**Figs. 2D and E**). In DksA itself, binding of ppGpp to the binary complex resulted in movements of DksA that include shifting of the α_1_, α_4_, and CT-helix toward the center of the G domain, swinging of α_2_, and α_3_ ahelices in the opposite direction, and slight bending of the CC domain, returning DksA to a conformation that more closely resembles that of unbound DksA (**Figs. 2G and H, Movie S3**). The electron density map around the G domain of DksA is well defined in the ternary complex (**Fig. S1E**) suggesting that the ppGpp binding enhances the DksA affinity to RNAP (**Fig. S1E**). These observations suggest that ppGpp binding relieves mechanical stress introduced by binding of DksA alone. The repositioning of DksA also explains the synergy between DksA and ppGpp in transcriptional inhibition and destabilization of RPo because when ppGpp is present, the CC tip is in a location that should severely compromise RNAP function.

### Structure of the RNAP–TraR complex

Although TraR appears to be a structural and functional homologue of DksA, ppGpp does not influence its activity (Blankschien et al., 2009b; Gopalkrishnan et al., 2017; Grace et al., 2015). To extend our understanding of the mechanism of transcriptional regulation through the secondary channel, we prepared crystals of the RNAP–TraR complex by soaking TraR into RNAP crystals and determined the structure at 3.8 Å resolution (**Table 1**). Strong electron density corresponding to TraR was observed from the βlobe/i4 domain extending through the secondary channel to the active site (**Figs. S1F and S2D**), allowing us to build the TraR model (**Fig. S1B**) and refine the RNAP–TraR complex structure (**Fig. 4A, Movie S6**).

**Figure 4.**
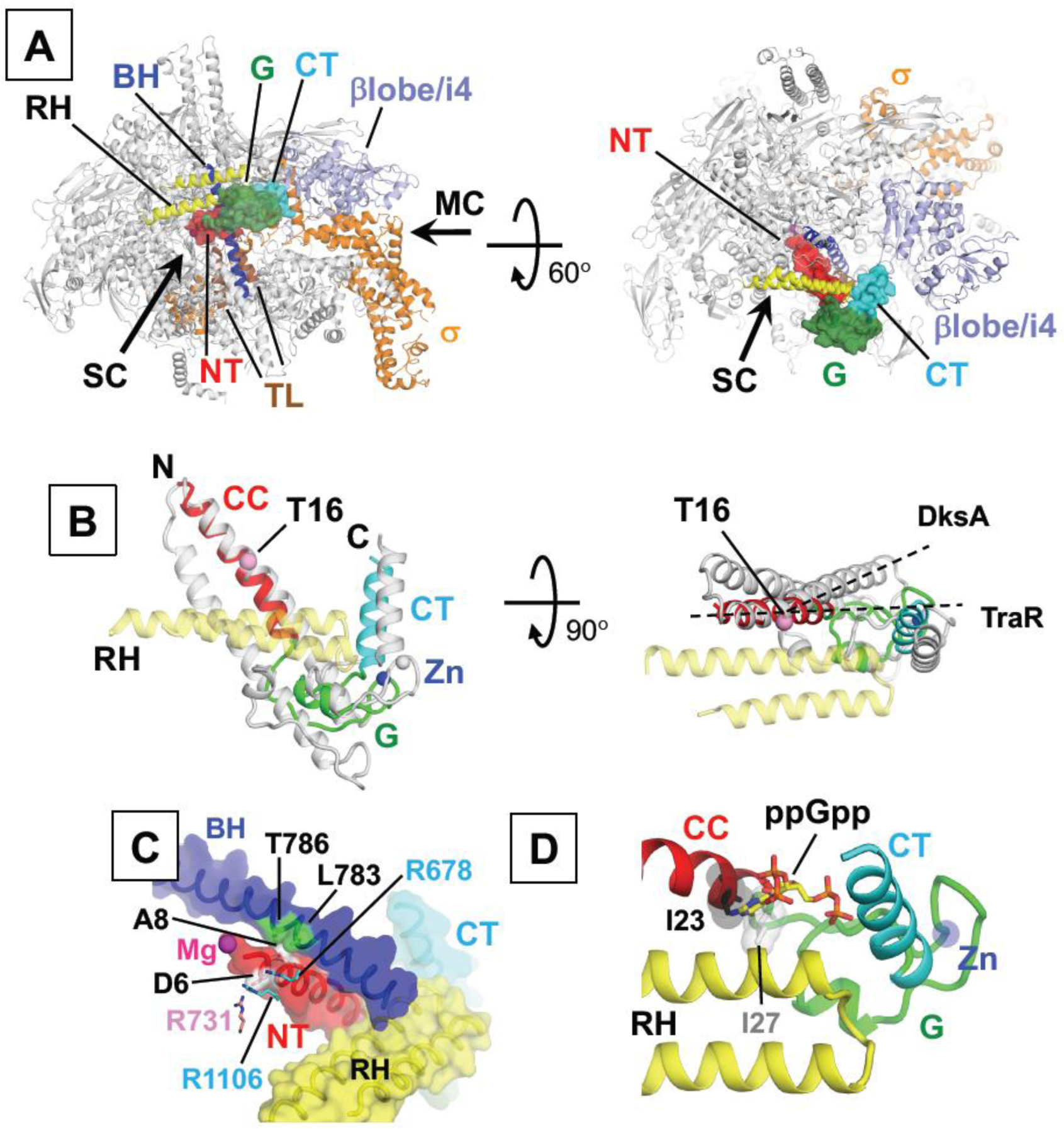
The RNAP - TraR complex. (**A**) RNAP is depicted as ribbons and TraR is depicted as a ribbon with a partially transparent surface. The relative orientations of the molecules are indicated in each panel. (**B**) Comparison of the RNAP–bound TraR (color image) and DksA (white). Positions of the N and C-terminus of TraR are indicated. The β’rim helix of RNAP is depicted as a partially transparent yellow ribbon model. Residue T16 of TraR (pink sphere) and the trajectories of α helixes of TraR and DksA interacting with the rim helix are depicted as black dashed lines. (**C**) A magnified view of the tip of the TraR NT-helix. TraR, BH and RH of RNAP are depicted as ribbons with partially transparent surfaces. The D6 and A8 residues of TraR and amino acid residues of RNAP contacting with these residues are depicted as sticks and labeled. (**D**) ppGpp binding site 2 is occupied by hydrophobic residues of TraR (I23 and I27).

The N-terminal α1 helix (NT-helix) of TraR, which is analogous to the second helix of the DksA CC domain, fits into the secondary channel of RNAP (**Fig. 4B**). Electron density of the TraR N-terminus is traceable beginning with the glutamate at position 4. As observed in the RNAP—DksA/ppGpp ternary complex, the D6 residue of TraR, which corresponds to D74 of DksA, is positioned near the NTP binding site (*i*+1 site) and may form salt bridges with R678/R1106 (β subunit) and R731 (β’ subunit). The A8 residue of TraR, the counterpart of A76 of DksA, faces the center of the BH and fits into the same cavity in the BH occupied by DksA A76 (**Fig. 4C**). Mutations of either of these amino acids in TraR, D6A or A8T, severely impaired TraR function (Blankschien et al., 2009b; Gopalkrishnan et al., 2017) demonstrating the importance of both interactions for TraR activity.

The NT-helix of TraR extends from the RNAP active site to the outer rim of the secondary channel where the G domain, which contains four Cys residues that coordinate a Zn atom, contacts the β’rim helix. The NT-helix initially follows the same path as the CC domain of DksA when bound in the presence of ppGpp (**Fig. 4B**). However, the trajectory of the NT-helix diverges from that of the DksA CC domain after TraR residue T16. This conformation brings the NT-helix and G domain of TraR into close proximity with the β’rim helix increasing direct interactions between TraR and RNAP. The hydrophobic residues I23 and I27 of TraR occupy the space filled by ppGpp clearly demonstrating why ppGpp does not affect TraR activity (**Fig. 4D**). The CT-helix of TraR extends away from the G domain and its C-terminus contact with the βlobe/i4 domain as observed in the RNAP and DksA complex. Unlike DksA alone, TraR binding does not trigger any major conformational change in RNAP. As a result, the RNAP–TraR complex is more similar to the RNAP—DksA/ppGpp complex than the RNAP—DksA complex. Like DksA, TraR is able to simultaneously access two important sites when bound to RNAP, the BH and one of the pincers that forms the DNA binding channel (the βlobe/i4 domain) indicative of their similar transcriptional regulatory activities.

### NTP entry into the RNAP active site in the presence of secondary channel binding transcription factors

The CC domain of DksA and the NT-helix of TraR fully occupy the secondary channel of RNAP, which has been proposed to serve as the major access route of NTPs to the active site (**Fig. S4A, B and C)** (Batada et al., 2004; Zhang et al., 2015). This fact raises the question of how NTPs can access the active site of RNAP in the presence of bound DksA or TraR. We therefore explored additional possible NTP loading pathways in RPo containing DksA or TraR by identifying empty spaces in these complexes available for NTP loading (**Fig. S4E**). Our modeling revealed that NTP can be loaded via the main channel because only the single stranded DNA is located in the main channel in RPo (**Fig. S4D**) as opposed to a ~9 bp long DNA/RNA hybrid in the elongation complex. Because the NTP loading path through the secondary channel is completely blocked by DksA or TraR, the main channel should become the sole pathway for NTPs to access the active site during transcript initiation in the presence of these factors (**Fig. S4E**).

### Determination of the affinities of DksA and TraR to the RNAP and the effect of ppGpp

A comparison of the structures of binary and ternary complexes suggests that the synergism observed between DksA and ppGpp could arise from ppGpp increasing the extent of the interaction surface between DksA and RNAP and eliminating the energetically unfavorable strained conformations of DksA and RNAP, activities that should increase the affinity of DksA for RNAP. To test this model, we measured the dissociation constant (Kd) of DksA with core enzyme and the σ^70^ holoenzyme using a fluorescence anisotropy assay (**Fig. 5**). A DksA derivative with a cysteine substitution at position 35 (A35C) was constructed and used to conjugate a fluorescent label to DksA. This site was chosen for labeling because it is surface exposed and located in a region that is not involved in binding to RNAP. Affinity was measured by adding increasing amounts of RNAP to labeled DksA (DksA^fl^) and measuring changes in anisotropy. DksA binds with slightly lower affinity to the core enzyme compared to the σ^70^ holoenzyme (K_d_=112 and 52 nM for core and holoenzymes, respectively) (**Fig. 5A**). When ppGpp was added to the binding reaction, the affinity of DksA to both the core and holoenzymes increased in a concentration-dependent manner (**Figs. 5C and D**).

**Figure 5.**
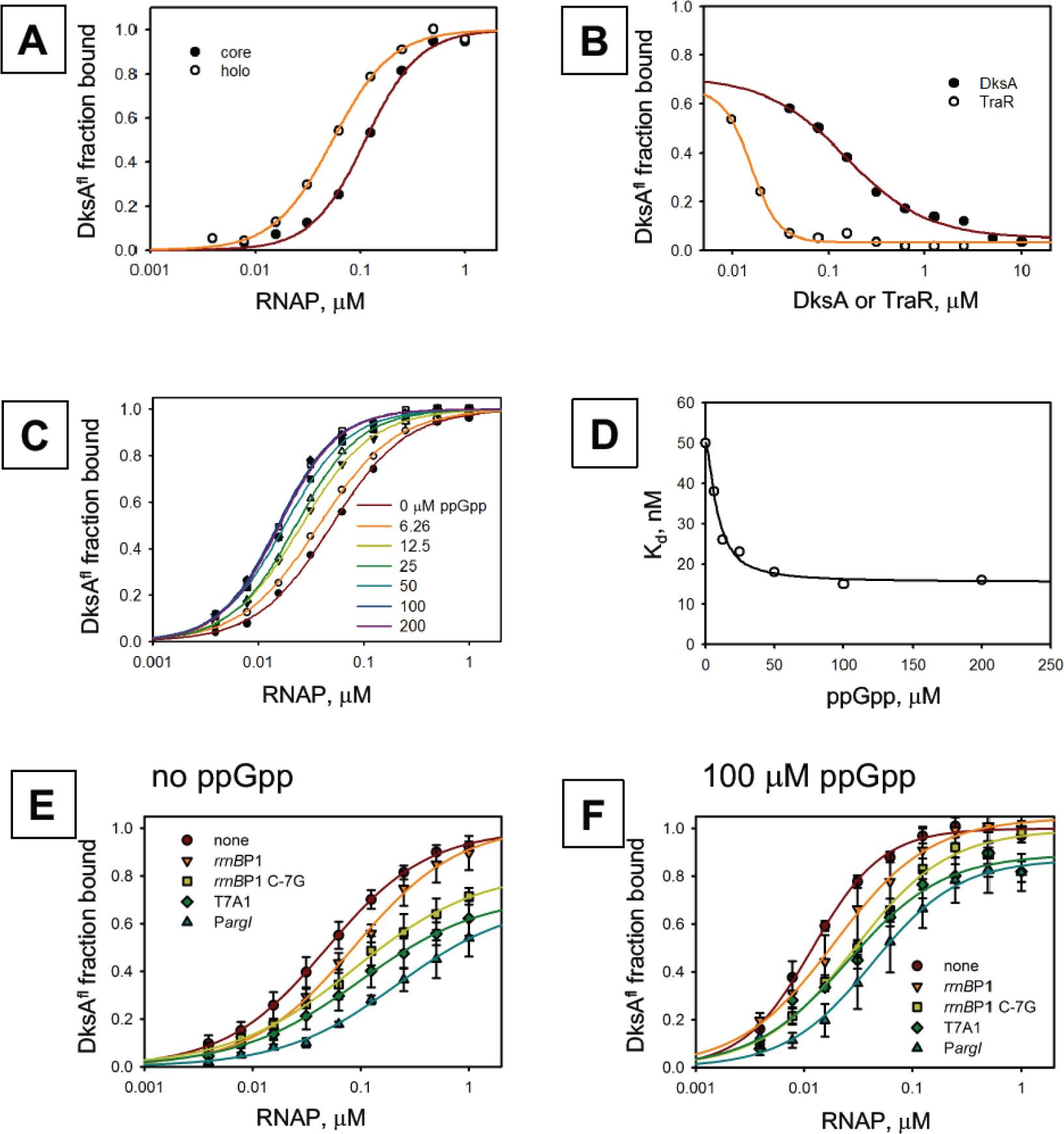
Apparent affinity of DksA with various forms of RNAP. Binding was measured using a fluorescence anisotropy assays with BODIPY FL-DksAo35A (DksA^fl^). (**A**) Binding to core RNAP and the s70 holoenzyme is shown. (**B**) A competition assay in which unlabeled DksA and TraR displace DksAfl is shown. TraR is able to compete for binding at lower concentrations than DksA indicating that the affinity of TraR for RNAP is greater than that of DksA. (**C**) Addition of increasing concentrations of ppGpp to the binding assay increases the apparent affinity of DksA for RNAP. (**D**) The concentration of DksA needed for half-maximal binding is shown for each concentration of ppGpp tested. Colors of the points correspond to the ppGpp concentrations indicated in panel C. (E-F) DksA binding is reduced for RNAP - promoter complexes compared to RNAP alone (red circles) in the absence (**E**) and presence of ppGpp (**F**), and ppGpp enhances the apparent affinity of DksA for the RNAP–DNA complexes as it does with RNAP alone.

The structure of TraR bound to RNAP indicates that it makes more extensive contacts with RNAP and therefore should bind with higher affinity than DksA. To test this hypothesis, we used a competitive binding assay in which the anisotropy of DksA^fl^ bound to RNAP was measured in the presence of increasing amounts of unlabeled TraR or unlabeled DksA. TraR was able to displace DksA^fl^ at significantly lower concentrations than DksA (**Fig. 5B**). Binding constants estimated from the competition experiment indicate that the affinity of TraR with RNAP is 6 nM, which is 8.6 and 3 times higher than the affinity of DksA with RNAP in the absence and presence of 100 μM ppGpp, respectively. We also note that the affinity of unlabeled DksA for RNAP measured using the competition experiment is 52 nM, which is comparable to that measured for DksA^fl^ indicating that the fluorescent label does not interfere with binding affinity.

To measure the binding of DksA^fl^ to DNA-bound RNAP, DksA was incubated with RNAP bound to several different promoter DNA fragments including the positively regulated *PargI* promoter, insensitive T7A1 promoter, and negatively regulated *rrnB*P1 promoter (**Figs. 5E and F, Fig. S5**) (Gummesson et al., 2013; Lyzen et al., 2009; Paul et al., 2004; Perederina et al., 2004). Promoter DNA was in 6-fold molar excess compared to RNAP to minimize the amount of free RNAP. The apparent affinity of DksA with the RNAP–promoter DNA complex was weaker than that for free holoenzyme for all promoters tested (**Figs. 5E**). There is an interesting correlation between the apparent DksA affinity and promoter type, *e.g.,* RNAP with the negatively regulated *rrnB*P1 promoter showed the highest affinity for DksA, followed by the insensitive T7A1 promoter. RNAP bound to the positively regulated P*argI* promoter had the weakest apparent affinity for DksA. To test the correlation between the type of regulation and apparent affinity of the RNAP–DNA complex for DksA, we introduced the C-7G mutation into the *rrnB*P1 promoter. This single base substitution dramatically increases the half-life of RPo and renders the promoter insensitive to DksA and ppGpp (Haugen et al., 2006). Interestingly, this mutation also reduced the apparent affinity of DksA for the RNAP–DNA complex. Our results are consistent with those obtained using an iron-mediated cleavage assay to detect DksA binding to RNAP alone and with the consensus “full con” promoter that forms a very stable RPo.Those experiments showed that DksA binding to RPo was reduced by approximately 10-fold compared to apo-form RNAP (Lennon et al., 2009).

We also tested the impact ppGpp on DksA binding to RNAP–DNA complexes by first incubating RNAP with the promoter DNA fragments, then adding DksA^fl^ and ppGpp to the RNAP–DNA complexes. Anisotropy of DksA^fl^ was measured after 5 min incubation. ppGpp increased the extent of DksA binding to all promoters (**Fig. 5F**). The overall pattern of apparent affinities was maintained with *rrnB*P1 exhibiting the highest affinity and P*arg*I the weakest of the promoter set.

## DISCUSSION

In this study, we present X-ray crystal structures of the *E. coli* RNAP holoenzyme in complex with DksA in the presence and absence of ppGpp (**Figs. 1, 2 and 3, Movies S1 and S4**). The structures show that DksA contacts two key areas of RNAP that are critical for maintaining the RPo, namely the BH, which positions the template DNA in the active site and the βlobe/i4 domain, which forms part of the clamp holding the downstream duplex DNA (**Figs. 1B and 2A**). The structures also suggest that the CC domain of DksA is positioned to potentially interfere with the catalytic step of RNA synthesis by preventing the folding of trigger loop and by binding to NTP at the active site (**Fig. S3**). The models of RNAP—DksA and RNAP—DksA/ppGpp complexes were established based on the cross-linking and mutagenesis studies (Furman et al., 2013; Parshin et al., 2015; Ross et al., 2016) and our crystal structures prepared by soaking the DksA into the *E. coli* RNAP holoenzyme crystals are in good agreement with these models.

ppGpp was discovered nearly 50 years ago as a key regulator of rRNA transcription (Cashel and Gallant, 1969). However, elucidation of the mechanism of regulation proved challenging because effects of ppGpp observed *in vitro* could not fully account for the magnitude of the effects seen *in vivo.* The discovery of DksA and findings demonstrating that DksA potentiated the effects of ppGpp significantly advanced our understanding of the regulatory mechanisms involved (Paul et al., 2004; Perederina et al., 2004). It has been proposed that DksA enhances the regulatory function of ppGpp by acting as a cofactor (Haugen et al., 2008; Hauryliuk et al., 2015; Potrykus and Cashel, 2008). The structure of DksA bound to RNAP reported here reveals that DksA binds to RNAP in a stressed conformation that involves distortions in both RNAP and DksA. ppGpp binding repositions DksA and restores both RNAP and DksA conformations to their original states (**Fig. 2, Movies S3 and S5**). Our structural and biochemical data indicate that ppGpp enhances DksA function by both increasing its affinity for RNAP (**Fig. 5C**) and by guiding the CC tip into a position that will effectively conflict with RPo formation and impinge on the catalytic step of RNA synthesis (**Figs. 2A and B**). Of the two ppGpp binding sites on RNAP, ppGpp binding at site 2, which is formed only in the presence of DksA, appears to be the key player in mediating the effects of ppGpp in transcription initiation (**Fig. 3**) (Ross et al., 2016). We therefore propose to re-evaluate the roles of DksA and ppGpp in their synergistic action on rRNA transcription inhibition via site 2 as follows: DksA possesses the ability to influence RNAP activity and ppGpp allosterically potentiates the activity of DksA. This model is supported by studies of the structure and function of TraR, including this study, showing that TraR possesses the same structural organization and mode of binding to RNAP as DksA, but influences RNAP transcription without ppGpp.

### Structural basis for transcription inhibition by DksA/ppGpp and TraR

From the structural analyses of the RNAP—DksA and RNAP—DksA-ppGpp complexes, we found that the CC tip of DksA is positioned near the active site of RNAP, consistent with previous biochemical studies probing the RNAP—DksA complex (Lennon et al., 2009; Parshin et al., 2015). In addition, ppGpp binding at the interface between RNAP and DksA inserts the CC tip deeper of DksA to the active site of RNAP, strengthening interactions between the CC tip and the BH (**Figs. 2A and B**). It has been proposed that although the BH is distantly positioned from the DNA binding clamp of RNAP, the interaction between the CC tip and the BH affects the conformation of the DNA binding clamp by an allosteric mechanism. As a result, the equilibrium among the several forms RNAP–promoter complexes shifts toward the less stable early-stage species (Rutherford et al., 2009). In the RPo, the template DNA lands on the BH and the interaction between a DNA base at *i*+1 site (+2 DNA base in the case of RPo) and T790 residue of the BH stabilizes the RPo (**Fig. 6**) (Zhang et al., 2012). Therefore, in addition to the allosteric mechanism, we propose that the physical contact between the CC tip and BH disfavors the interaction between the BH and the template DNA and accordingly destabilizes the RPo.

**Figure 6.**
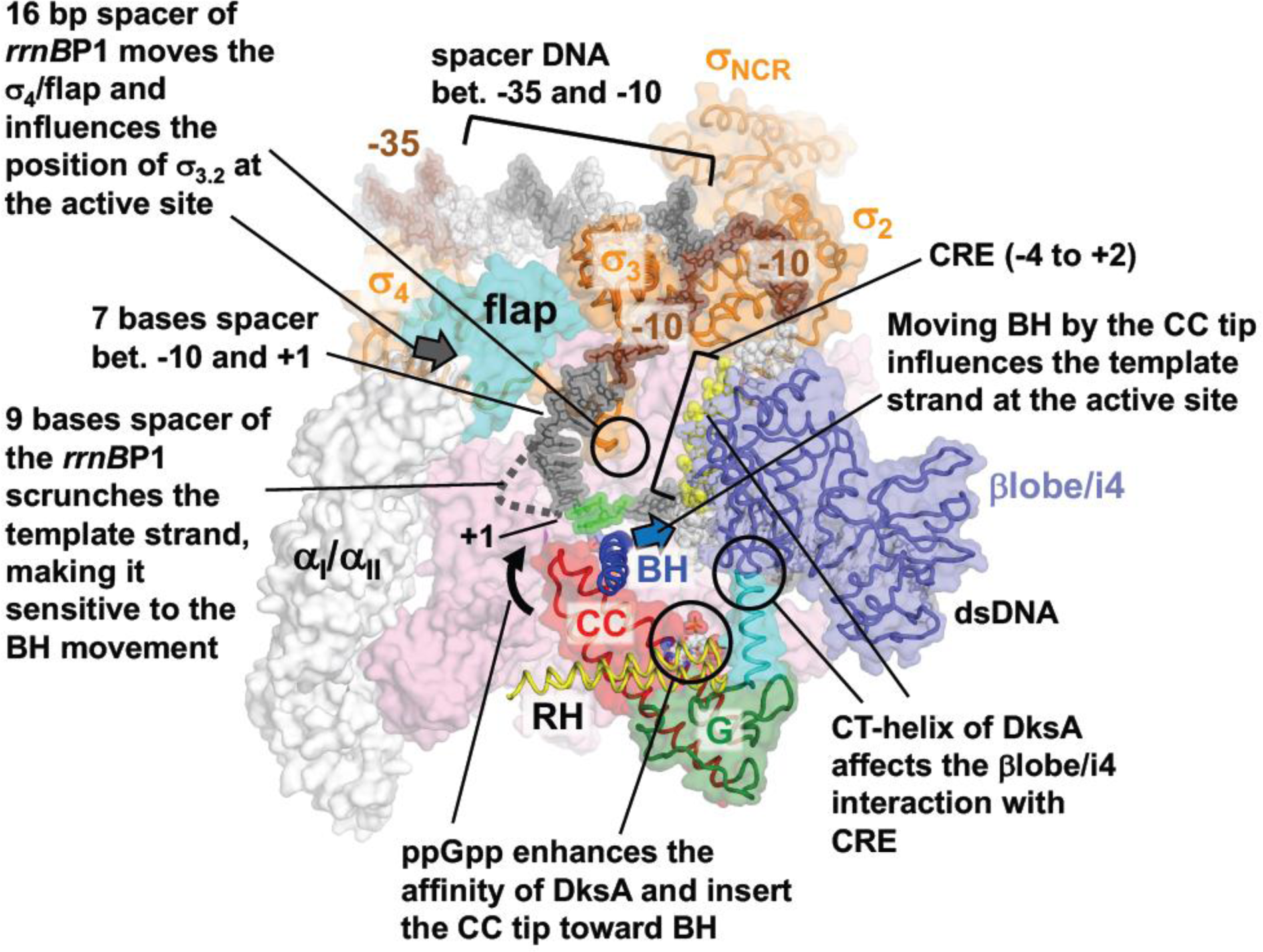
A model for the structural basis of *rrnB*P1 transcription inhibition by DksA/ppGpp. A model of RPo with DksA and ppGpp was constructed by superposing the structures of the RNAP-DksA/ppGpp ternary complex and the RNAP transcription initiation complex (PDB: 4YLN). Template and non-template DNA strands are depicted in white and black CPK representation with -35/-10 elements (brown), core recognition element (CRE, yellow) and the transcription start site (+1 and +2, green). Domains and motifs of RNAP and DksA playing key roles in transcription inhibition are labeled. For clarity, β subunit is removed except the flap and i4/lobe domains. There are 7 bases between the -10 element and the transcription start site (+1) in this model, suggesting the DNA is scrunched in the open complex formed with *rrnB*P1 promoter.

This structural model for transcription inhibition is supported by many biochemical studies with the *rrnB*P1 promoter. This promoter is one of the strongest promoters in the *E. coli* genome, and it has been investigated as a model promoter for understanding transcriptional regulation in response of changing growth conditions. The suboptimal organization of the *rrnB*P1 promoter, including the short 16 bp spacer between the -35 and -10 promoter elements, the GC-rich discriminator sequence, and the location of the transcription start site positioned 9 bases downstream from the end of the -10 element (Winkelman et al., 2015), results in formation of inherently unstable RPo. The extra bases between the -10 element and the transcription start site require additional scrunching of the template DNA in the RPo (**Fig. 6**), which creates tension likely making the RPo more sensitive to movement of the BH resulting from its interaction with the CC tip of DksA. Consistently, a *rrnB*P1 derivative having the C-7G mutation that shifts the transcription start site 6 bases downstream from the -10 element thereby reducing scrunching forms a DksA/ppGpp-insensitive RPo (Haugen et al., 2006; Winkelman et al., 2015).

The spacer between the -35 and -10 promoter elements varies for σ^70^-dependent promoters with 17 bp being the most common. The spacer in *rrnB*P1 the promoter is 16 bp, and this difference also contributes to the sensitivity of *rrnB*P1 to negative regulation by ppGpp and DksA. To recognize promoters with different spacer lengths, the distance between σ domain 4 (σ_4_) and σ domain 2 (σ_2_) responsible for recognizing the -35 and -10 elements, respectively, must be adjusted (Zuo and Steitz, 2015). The RPo formed at a promoter with a 16-bp spacer, like *rrnB*P1, requires that σ_4_ shifts by ~4 Å, equivalent to a 1-bp translocation in double-strand DNA, compared to RPo at a promoter with a 17-bp spacer. The motion of σ_4_ within RNAP is likely linked to the adjacent σ region 3.2 (σ_3.2_), which penetrates into RNAP active site and directly contacts the template strand helping to stabilize RPo (Zhang et al., 2012). Therefore, it is tempting to speculate that the short spacer of *rrnB*P1 influences the position of σ_3.2_ making this RPo more sensitive to changes in the position of the BH caused by its interaction with the CC tip of DksA (**Fig. 6**). Consistent with this model, increasing the *rrnB*P1 spacer to 17 bp also makes this promoter insensitive to DksA and ppGpp (Haugen et al., 2006).

The βlobe/i4 domain have been found to be critical for DksA binding to RNAP (Parshin et al., 2015). The structural studies presented here confirm that direct contacts are made between DksA and this domain of RNAP. In addition, the structures suggest that interactions with the βlobe/i4 domain may also contribute to regulation beyond providing a docking site for DksA. The βlobe/i4 domain interacts with the core recognition element (CRE) in the non-template strand of promoter DNA from the -4 to +2 position. This interaction is proposed to contribute to the formation and maintenance of a stable transcription bubble in the RPo (**Fig. 6, Fig. S6A**) (Petushkov et al., 2015; Zhang et al., 2012). Contact between the CT-helix and the βlobe/i4 domain in the absence of ppGpp shifts the position of the βlobe/i4 domain relative to the main body of RNAP (**Fig. 2C, Movie S2**) in a way that may weaken its grip on the non-template DNA in the transcription bubble and consequently decrease RPo stability. The importance of the βlobe/i4 domain for the DksA-dependent transcriptional repression is further supported by the isolation of Δ*dksA* suppressor mutations in the βlobe domain and also around linkers connecting the lobe domain to the main body of RNAP (Rutherford et al., 2009) (**Fig. S6A**).

Some *ΔdksA* suppressor mutations have been mapped to the switch regions of RNAP that serve as hinges for the RNAP clamp and undergo conformational changes during the opening and closing of the main channel in the course of RPo formation (Feklistov et al., 2017; Srivastava et al., 2011). These suppressor mutations mimicked the negative effects of DksA on *rrnB*P1 transcription by inhibiting transcription and dramatically reducing RPo stability (Rutherford et al., 2009). In the crystal structures of RNAP determined in this study, the DNA binding channel is in the closed conformation and has to be opened to load DNA into the channel (Feklistov et al., 2017). Although the association of DksA and ppGpp with RNAP does not influence the conformation of the main channel (**Fig. 2E**), we speculate that the interaction between DksA and the βlobe/i4 domain may restrict opening of the channel in the course of the RPo formation thereby altering the dynamics in favor of the closed complex.

Of all the secondary channel binding proteins, the activities of DksA and TraR on transcription initiation are the most similar. Both factors decrease transcription of ribosomal genes and increase transcription of amino acid biosynthesis genes (Gopalkrishnan 2017) (Blankschien et al., 2009b). Despite the lack of sequence conservation, our results clearly show that the structures of DksA and TraR are remarkably similar (**Fig. 4B, Figs. S1A and B**). Although TraR lacks the first 68 amino acids DksA, which corresponds to part of the G domain and the entire first helix of the CC domain, these factors interact with RNAP in a similar manner (**Figs. 2B and 4C**) by reaching into the active site via the elongated CC domain/NT-helix and interacting with the DNA binding βlobe/i4 domain via the C-terminal CT-helix.

Binding of DksA to RNAP is similar to that observed for the Gre factors (**Fig. S7**) (Opalka et al., 2003; Sekine et al., 2015). Each of these proteins inserts an elongated CC domain into the secondary channel of RNAP and positions the CC tip near the active site. However, structural changes in RNAP caused by binding of GreA and GreB are different. The Gre factors bind and wrap around the edge of β’rim helix with the G domain but do not distort the structure of the β’rim helix (Sekine et al., 2015). In addition, the Gre factors lack a counterpart of the CT-helix of DksA (**Fig. S1A, B and C**) and therefore are unable to influence the stability of the RPo via interaction with the βlobe/i4 domain.

### Synergism between ppGpp and DksA during transcription inhibition

ppGpp increases the affinity of DksA for RNAP approximately 3-fold (**Fig. 5C**). In the absence of ppGpp, DksA binding to RNAP alters the conformations of the β’rim helix, the βlobe/i4 domain, and DksA introducing strain into the complex. In the presence of ppGpp, DksA is reoriented such that it binds along the rim helix in a way that does not cause conformational distortions in RNAP or DksA and increases the contact area between the two proteins (**Fig. 2, Movie S2**). The detrimental effects of the distortions in RNAP caused by DksA binding on DksA affinity for RNAP are further supported by the finding that TraR does not alter the conformation of RNAP and binds significantly tighter to RNAP than DksA (**Fig. 5B**).

Amino acid substitutions in DksA, such as L15F and N88I, bypass the requirement for ppGpp in *E. coli* cells growing in nutrient-depleted media. These DksA derivatives, known as super DksAs, increase the activity of the transcription factor and work independently of ppGpp (Blankschien et al., 2009a). Interestingly, these residues are not involved in binding to RNAP. Instead, they are located in the hydrophobic core of the G domain (L15) and in the center of the CC domain (N88); namely areas of DksA that undergo large conformational changes during the formation of the RNAP—DksA binary complex (**Movie S3),** These findings suggest that the amino acid substitutions may act by stabilizing a DksA conformation similar to that in the tertiary complex containing RNAP, DksA and ppGpp.

### ppGpp binding site 1

The DksA-independent ppGpp binding site 1 has been identified by crystallographic studies of *E. coli* RNAP in complex with ppGpp (Mechold et al., 2013; Zuo et al., 2013). Although the same ppGpp binding site was identified, different orientations of ppGpp were observed. Mechold *et al.* proposed that the 3’-diphosphate group of ppGpp is positioned near the β’ subunit, whereas Zuo *et al.* assigned the same electron density to the guanine base in accordance with cross-linking data (Ross et al., 2013). Experimental evidence compiled from structural and biochemical studies reveals the correct orientation of ppGpp at the site 1 as follows (**Fig. 3B**). The 5’ - and 3’-phosphate groups face the ω subunit due to their larger electron densities compared with a conformation in which these groups are oriented toward the β’ subunit. The 5’- and 3’-phosphate groups face the R3 and R52 residues of the ω subunit, respectively. This orientation is also supported by a comparison of the structures of RNAP with ppGpp and pppGpp (Mechold et al., 2013). The guanine base faces the β’ subunit, which is supported by the ^32^P-6-thio-ppGpp crosslinks to β’ subunit (Ross et al., 2013). In this orientation, His/Asp residues form hydrogen bonds with the guanine base, explaining the specificity for guanine binding at site 1.

### Activation of Transcription by ppGpp and DksA

The structures described in this work provide a straightforward model for how TraR and DksA alone and together with ppGpp negatively regulate transcription initiation. However, DksA/ppGpp and TraR are also able to activate transcription at promoters such as those encoding genes for amino acid biosynthesis (Gopalkrishnan et al., 2017; Paul et al., 2005) and promoters regulated by the alternative sigma factor σ^E^ (Costanzo et al., 2008; Gopalkrishnan et al., 2014; Grace et al., 2015). Mechanisms of transcription activation via these secondary channel binding proteins cannot be elucidated directly from the crystal structures determined in this study. Differences between the structures of DksA with and without ppGpp reveal the flexible nature of DksA’s interaction with RNAP. Promoters that are positively regulated by DksA and ppGpp tend to form stable RPo suggesting that the mechanism of action involves distinct interactions with RNAP as compared to inhibition. Interestingly, DksA and ppGpp only activate and do not inhibit transcription by the holoenzyme containing σ^E^, which is a member of the group 4 class of σ factors that are defined by a reduced domain structure as compared to the primary σ factor. σ^E^ and other group 4 sigma factors have domains 2 and 4, which are responsible for binding the -35 and -10 promoter elements, but lack most of domain 1 including region 1.1 and have a short linker in place of domain 3. The differences in domain structure between σ^70^ and σ^E^ may play a role in restricting regulation by DksA and ppGpp to activation. Biochemical and structural studies with the σ^E^ holoenzyme in the absence and in the presence of DksA/ppGpp or TraR may better elucidate the basis for the activation since an inhibitory state is not accessible in this transcription system.

## EXPERIMENTAL PROCEDURES

### Purification of σ^70^ RNAP holoenzymes

Expression, purification and reconstitution of wild-type and mutant *E. coli* σ^70^ RNAP holoenzymes were performed as previously described (Molodtsov et al., 2016).

### Purification of DksA

A 3L culture containing BL21(DE3) cells transformed with a pSUMO-DksA vector, which contains the *E. coli dksA* gene cloned to express a His_6_-SUMO-DksA fusion, was grown at 30 °C in LB medium supplemented with 50 μg/ml Kanamycin and 200 μM ZnCl_2_, induced with 1 mM IPTG at OD_600_ = 0.7, and incubated for an additional 3 hours to allow for protein expression. Cells were harvested by centrifugation and lysed by sonication in lysis buffer (20 mM Tris-HCl, pH 8.0 at 4°C, 500 mM NaCl, 5% glycerol, 0.1 mM EDTA, 2 mM PMSF). The lysate was clarified by centrifugation for 30 min at 17,000 g, 4°C, and the supernatant was mixed with 1.5 ml of Ni agarose slurry (Qiagen) equilibrated with lysis buffer and incubated on a shaker for 2 hours at 4 °C. The Ni resin was washed first with 30 column volumes of lysis buffer and then with 30 column volumes of wash buffer (10 mM Tris-HCl, pH 8.0 at 4°C, 150 mM NaCl, 0.1 mM EDTA) supplemented with 20 mM imidazole. Elution was carried using wash buffer supplemented with 300 mM imidazole. The eluted protein was diluted with wash buffer lacking NaCl to reduce the salt concentration to 70 mM and applied to a 1 ml sepharose Q column (GE Healthcare) equilibrated with the same buffer. SUMO-DksA was eluted by a linear gradient of 0.07-0.5 M NaCl over 40 column volumes. Fractions containing SUMO-DksA were diluted with wash buffer lacking NaCl to reduce the salt concentration to 70 mM and incubated with UlpI protease to remove the SUMO tag overnight at 4°C. The sample was next mixed with 1.5 ml of equilibrated Ni agarose slurry (Qiagen), incubated on a shaker for 2 hours at 4°C, and centrifuged for 5 min at 6,000g at 4°C. The supernatant containing wild-type DksA lacking the His tag was analyzed for purity by SDS-PAGE and concentrated using VivaSpin concentrators. Aliquots of 1 mM DksA were frozen in liquid N2 and stored at - 80°C until soaking to crystals.

### Purification of TraR

A 3L cell culture of BL21(DE3) *E. coli* cells carrying pET24a-TraR-His6, pET24a vector with *E.coli traR* gene having a hexahistidine tag at the C-terminus (Blankschien et al., 2009b), was grown in LB medium at 30°C supplemented with 200 μM ZnCl_2_, induced with 1 mM IPTG at OD_600_ = 0.7, and incubated for an additional 3 hours to allow for protein expression. The cells were harvested by centrifugation and lysed by sonication in lysis buffer. The lysate was clarified by centrifugation at 4°C for 30 min at 17,000g. The supernatant was mixed with 1.5 ml of Ni agarose slurry (Qiagen) equilibrated with lysis buffer and incubated on a shaker for 2 hours at 4°C. The Ni resin was washed with 30 column volumes of the lysis buffer followed by 30 column volumes wash of wash buffer supplemented with 20 mM imidazole. Elution was carried using the wash buffer supplemented with 300 mM imidazole. Eluted proteins were diluted with the wash buffer lacking NaCl to reduce the salt concentration to 70 mM and applied to a 1 ml sepharose Q column (GE Healthcare) equilibrated with the same buffer. TraR was eluted by a linear gradient of 0.07-0.5 M NaCl over 40 column volumes. Fractions containing TraR were analyzed by SDS-PAGE and concentrated using VivaSpin concentrators. Aliquots of 700 μM TraR were frozen in liquid N2 and stored at - 80°C until soaking to crystals.

### Crystallization of σ^70^ holoenzyme-DksA and TraR complexes

Crystallization of the *E. coli* σ_70_ RNAP holoenzyme was performed as previously described (Murakami, 2013). To form co-crystals of RNAP and DksA or TraR, holoenzyme crystals were transferred to the cryoprotection solution (0.1 M HEPES (pH 7.0), 0.2 M calcium acetate, 25 % PEG400, 10 mM DTT) supplemented with 0.2 mM DksA or TraR and incubated overnight at 22 °C followed by flash-freezing in liquid N_2_. To form crystals of RNAP–DksA/ppGpp, crystals soaked overnight with DksA were transferred to a fresh cryoprotection solution (0.1 M HEPES (pH 7.0), 0.02 M MgCl_2_, 25% PEG400, 10 mM DTT) supplemented with 1 mM ppGpp and 0.05 mM DksA, incubated for 2 hours, and then flash-frozen in liquid N_2_.

There are two RNAP molecules in an asymmetric unit of the RNAP crystal (**Fig. S8**). For crystals of RNAP—DksA and RNAP–TraR complexes, both RNAP molecules form complexes with DksA or TraR. In contrast, the crystal of RNAP—DksA/ppGpp complex showed that only one RNAP molecule is associated with DksA. Investigation of the crystal packing of the RNAP–DksA/ppGpp complex showed that the DksA binding site of the second RNAP molecule is blocked by the α subunit from the symmetry related RNAP. Binding between the RNAP (second molecule) and αCTD (from the first molecule) was triggered by replacement of the divalent cation in the crystallization solution from calcium acetate (for soaking DksA) to MgCl_2_ (for soaking DksA and ppGpp) due to poor solubility of ppGpp in the presence of calcium acetate. Binding between the second RNAP molecule and the αCTD from the first molecule of RNAP was also observed in the crystal structure of *E. coli* RNAP alone when MgCl_2_ was included in the crystallization solution (unpublished).

### X-ray crystallographic data collection

The crystallographic datasets were collected at the Macromolecular Diffraction at the Cornell High Energy Synchrotron Source (MacCHESS) (Cornell University, Ithaca, NY) and the data were processed by HKL2000 (Otwinowski and Minor, 1997). The resolution limits for crystallographic datasets were determined based on completeness (>80 %) and CC_1/2_ (>20 %) rather than R_merge_ and <I>/σI > 2 criteria, since this approach prevents loss of useful crystallographic data for structure refinement as found in a recent study (Karplus and Diederichs, 2012). The structures were solved by molecular replacement using the suite of programs PHENIX (Afonine et al., 2010) and *E. coli* RNAP holoenzyme (PDB: 4YG2) (Molodtsov et al., 2015) as a search model. Strong Fo-Fc maps corresponding to DksA, ppGpp and TraR were observed after the rigid body refinement. A crystal structure of DksA (Perederina et al., 2004), a homology model of TraR and ppGpp were fitted into these extra density maps to continue the refinement. The program Coot (Emsley and Cowtan, 2004) was used for manual adjustment of the models during refinement. A homology model of TraR was constructed by SWISS-MODEL (Biasini et al., 2014) using DksA (Perederina et al., 2004) and Zn-finger binding protein YBIL from *E. coli* (PDB: 2KGO) as reference structures. The structures were refined by using the Phenix suite of programs for the rigid body and positional refinements with non-crystallographic symmetry and reference structure restraints to avoid over-fitting the data (R_free_ – R_work_ is less than 6 %), and to maintain the Ramachandran outliers less than 2 %. B-factors were refined as group B-factors since the data to parameter ratio is low. Final coordinates and structure factors were submitted to the PDB depository with ID codes listed in **Table 1**.

### Fluorescence anisotropy assay

N-terminally His-tagged DksA-C35A was purified as previously described (Costanzo Mol Micro 2008) and stored in storage buffer (20 mM Tris-HCl (pH 8), 150 mM NaCl, 1 mM ZnCl_2_ 10 mM 2-mercaptoethanol, 15% glycerol). Immediately before labeling, the storage buffer was exchanged via a Bio-Gel P4 desalting column (Bio-Rad) equilibrated with a solution containing 50 mM HEPES (pH 7.4), 500 mM NaCl, 1 mM ZnCl_2_, and 10% glycerol. The protein solution (10 μM) was then incubated on ice with sub-equimolar concentration of BODIPY-FL maleimide (Invitrogen/Molecular Probes) diluted from a 2.5 mM BODIPY-FL stock in DMSO. After 45 min of incubation, the reaction was stopped by the addition of 10 mM 2-mercaptoethanol. Unreacted dye was removed by passing the sample twice through a Bio-Gel P4 desalting column equilibrated in a solution containing 50 mM Tris-HCl (pH 8), 500 mM NaCl, 1 mM ZnCl_2_, and 10% glycerol. The degree of dye labeling was determined by measuring the absorbance of the labeled protein at 505 nm (ε505 = 79,000 M^−1^ cm^−1^; ε280 = 1,300 M^−1^ cm^−1^) and was typically > 50 %.

Double stranded promoter DNAs were annealed from corresponding oligonucleotides (**Fig. S7**). RNAP–promoter DNA complexes were assembled by incubation of 2 μM RNAP and with 12.5 μM of the corresponding DNA construct in in transcription buffer (50 mM Tris-HCl (pH 8), 10 mM MgCh, 50 mM NaCl, 5% glycerol, 10 mM 2-mercaptoethanol, 0.01% NP-40).

Binding reaction mixtures (20 μl) containing 5 nM BODIPY FL-DksA_C35A_ and 0 to 200 μM ppGpp and 0 to 1 μM RNAP (or RNAP–DNA complexes) in transcription buffer (50 mM Tris-HCl (pH 8), 10 mM MgCl_2_, 50 mM NaCl, 5% glycerol, 10 mM 2-mercaptoethanol, 0.01% NP-40) were incubated at 37°C for 1 h for RNAP and 5 min for RNAP–DNA. Anisotropy was measured on an Infinite M1000 multimode plate reader from Tecan instruments at 30 °C (excitation of 470 nm and emission of 514 nm). Fractional occupancies were calculated as follows

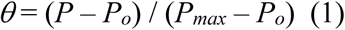

where *P*_0_ and *P* are polarization values before and after the addition of ligands, respectively, and *P*_max_ is the polarization value at saturation. Binding constants were obtained by fitting of the results to the Hill equation by non-linear regression using SigmaPlot.

Competition assays were used to determine the affinity of unlabeled DksA, TraR and DksA mutant derivatives for holo RNAP (Eσ^70^) and to determine the competition of DksA with promoter DNA. In these assays, reaction mixtures (20 μl contained BODIPY FL-DksA_C35A_, 100 nM RNAP, and either 0 or 50 μM ppGpp and 0 to 2 μM of unlabeled competitor proteins (e.g. TraR or DksA mutant) or promoter DNA duplexes. Components were incubated for 1 h at 37°C and fluorescence anisotropy was then measured. The 50% inhibitory concentrations (IC50) were calculated by using the Hill equation, and equilibrium dissociation constants (*K_I_*) were calculated from the IC_50_ as follows

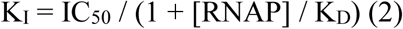

where *K_D_* refers to the affinity of BODIPY FL-DksA_C35A_ to RNAP (Cheng and Prusoff, 1973).

### Analysis of channels for NTP substrate entry based on crystal structures of RNAP

To search for potential pathways for NTPs to enter the active site of RNAP, we adopted the program CAVER (Chovancova et al., 2012) to identify cavities in four static structures including the apo-form holoenzyme (PDB: 4YG2), RNAP—DksA binary complex, RNAP–DksA/ppGpp ternary complex and RNAP–TraR complex. We note that all structures lack σ region 1.1 (σ_1.1_), which was shown to be located outside of the main channel when RNAP forms the open complex (Mekler et al., 2002). Therefore, it can be omitted from the NTP substrate entry analysis. In our analysis, the NTP molecule was modeled as a sphere of radius 3.5 Å as done in previous studies (Batada et al., 2004; Zhang et al., 2015), and default input parameters were used in CAVER3.01 except for shell_radius = 30 Å and shell_depth = 50 Å (see (Chovancova et al., 2012) for details). The NTP model sphere of radius 3.5 Å was used intially to search for potential pathways of entry. Once pathways had been identified, we used the whole molecule (in the all-atom representation instead of sphere) to further verify that NTPs could be accommodated (see below). For each structure, the averaged coordinates of the template DNA at +2 and +3 positions were chosen as the starting point for the pathway search in the main channel, while those of magnesium ion and +2 DNA were utilized for the secondary channel. We then identified the most probable pathway of NTP entry in both channels according to the available empty space and pathway distance (see (Chovancova et al., 2012) for methodology details).

### Molecular models of NTP along the substrate entry pathways

We applied k-center clustering on all the points that constitute the pathways and divided them into 10 microstates according to their positions. The geometric center of each microstate was extracted as the reference position for modelling NTP along the pathways. CTP coordinated with one magnesium ion was chosen to pair with the template DNA at +2 site. In particular, we aligned the center of mass of CTP with Mg^2+^ to each microstate center followed by combine with the specific RNAP holoenzyme structure to build the model. We then solvated the whole complex in a dodecahedron water box and added enough sodium ions to neutralize the system. The size of the box was defined with box edges 1 Å away from the protein surface. We first froze protein and performed 5,000-step energy minimizations on promoter DNA and NTP to let them re-orient to avoid the spatial clash. Second, we put positional restrain on all the heavy atoms of protein (with a force constant of 10 kJ•mol^−1^• Å ^−2^) and carried out another 10,000-steps energy minimization for nucleic and NTP to make them further fit to their protein surroundings. Finally, the whole system was energy minimized for 10,000 steps to achieve the molecular model of NTP along the pathway. All energy minimizations were performed with Gromacs4.5 (Pronk et al., 2013). Amber99sb force field (Hornak et al., 2006) were used for the whole system with parameter modifications on the polyphosphate tail (Meagher et al., 2003) of NTP molecule.

## AUTHOR CONTRIBUTIONS

V.M., S.E.A. and K.S.M. contributed to the design of the experiments and wrote the manuscript. V.M. prepared the crystals of RNAP in complex with DksA, DksA/ppGpp and TraR and collected the X-ray diffraction data, and K.S.M. determined their X-ray crystal structures. M.C. prepared ppGpp. L.Z. and X.H. analyzed the structures of RNAP to identify the NTP entry pathways to the active site. E.S. carried out the binding assays and analyzed the data with S.E.A. All authors discussed the results and commented on the manuscript.

## ACKNOWLEDGEMENTS

We thank the staff at the CHESS/MacCHESS at Cornell University for support of crystallographic data collection. We thank Christophe Herman for providing the expression vector of TraR and thank Shoko Murakami for constructing pSUMO–DksA vector. We thank Andrey Kulbachinskiy, Irina Artsimovitch, Nikolay Zenkin, Sergei Borukhov, Vicky Shingler, Richard Gourse, Wilma Ross and Kasia Potrykus for critically reading the manuscript. This work was supported by NIH grants GM087350 (K.S.M.) and GM097365 (S.A.), and Hong Kong Research Grants Council HKUST C6009-15G (X.H.), and also funded in part by the Intramural Research Program, NIH (M.C.).

---

**Table 1:** Data collection and refinement statistics

| RNAP complex with<br>PDB code | **DksA**<br>5W1T | **DksA/ppGpp**<br>5VSW | **TraR**<br>5W1S |
| --- | --- | --- | --- |
| **Data collection** |  |  |  |
| Space group | P2<sub>1</sub>2<sub>1</sub>2<sub>1</sub> | P2<sub>1</sub>2<sub>1</sub>2<sub>1</sub> | P2<sub>1</sub>2<sub>1</sub>2<sub>1</sub> |
| Cell dimensions |  |  |  |
| *a* (Å) | 184.0 | 187.3 | 186.5 |
| *b* (Å) | 204.9 | 205.3 | 206.0 |
| *c* (Å) | 314.3 | 311.3 | 310.3 |
| Resolution (Å) | 50 – 4.5 | 50 – 4.3 | 50 – 3.8 |
| Total reflections | 474,850 | 531,876 | 911,081 |
| Unique reflections | 70,723 | 78,563 | 117,447 |
| Redundancy | 6.7 (5.7) | 6.8 (6.4) | 7.8 (7.4) |
| Completeness (%) | 99.9 (99.9) | 95.6 (95.3) | 100.0 (100.0) |
| *I* / $\sigma$ | 16.6 (1.26) | 12.2 (1.06) | 17.5 (0.82) |
| Wilson B factor | 199.4 | 187.8 | 156.5 |
| *R*<sub>sym</sub> (%) | 10.0 (>100) | 12.8 (>100) | 9.2 (>100) |
| CC<sup>1/2</sup> | (0.360) | (0.373) | (0.270) |
| **Refinement** |  |  |  |
| Resolution (Å) | 50 – 4.5 | 50 – 4.3 | 50 – 3.8 |
| *R*<sub>work</sub> | 0.220 | 0.221 | 0.208 |
| *R*<sub>free</sub> | 0.276 | 0.259 | 0.262 |
| No. of atoms | 58,066 | 57,643 | 56,825 |
| *B* factors | 291.6 | 279.0 | 239.0 |
| R.m.s deviations |  |  |  |
| Bond length (Å) | 0.007 | 0.005 | 0.005 |
| Bond angles (°) | 1.147 | 1.08 | 1.068 |
| Clashscore | 16.8 | 17.2 | 17.6 |
| Ramachandran favored, % | 89.6 | 88.9 | 89.1 |
| Ramachandran outliers, % | 2.02 | 2.17 | 2.14 |

Data sets were collected at MacCHESS F1 line, Ithaca, NY
Highest resolution shells are shown in parentheses
Resolution limit of using the traditional criterion of $I/\sigma I > 2.0$ are: RNAP-DksA complex, 4.75 Å; RNAP-DksA/ppGpp complex, 4.73 Å; RNAP-TraR complex, 4.09 Å

**Supplemental Movie 1. Crystal structure of the *E. coli* RNAP and DksA binary complex.**

RNAP is depicted as ribbons and DksA is depicted as a ribbon with a partially transparent surface. Domains, motifs and regions of RNAP and DksA discussed in the text are labeled.

**Supplemental Movie 2. Binding of DksA to RNAP triggers conformational changes in RNAP**.

This movie shows conformational changes in RNAP that occur during formation of the binary complex (from the apo-forms of RNAP and DksA to the binary complex). RNAP and DksA are depicted as ribbons, and domains and regions of RNAP and DksA are colored as in Fig. 1A. The first scene shows a view from the RNAP main channel and the second scene shows a side view of a model of RNAP with promoter DNA and iNTP bound at the active site.

**Supplemental Movie 3. Conformational changes in DksA triggered by RNAP and ppGpp binding**.

This movie shows conformational changes in DksA that occur upon RNAP binding (first scene, from the apo-forms of RNAP and DksA to the binary complex) followed by ppGpp binding (second scene, from the binary to ternary complexes). The three structural domains of DksA and amino acid residues that are mutated in super DksA eliminating the need for ppGpp are indicated.

**Supplemental Movie 4. Crystal structure of the *E. coli* RNAP—DksA/ppGpp ternary complex.**

RNAP is depicted as ribbons and DksA is depicted as a ribbon with partially transparent surface. ppGpp molecules bound at the sites 1 and 2 are shown as CPK models. Domains, motifs and regions of RNAP and DksA discussed in the text are labeled.

**Supplemental Movie 5. ppGpp binding to the RNAP—DksA binary complex triggers conformational changes in RNAP and DksA**.

This movie shows conformational changes in RNAP and DksA that occur during the ternary complex formation (from the binary complex to the ternary complex). The first scene shows a view from the RNAP main channel and the second scene shows a side view of a model of RNAP with promoter DNA and iNTP bound at the active site.

**Supplemental Movie 6. Crystal structure of the *E. coli* RNAP and TraR binary complex.**

RNAP is depicted as ribbons and TraR is depicted as a ribbon with partially transparent surface. Domains, motifs and regions of RNAP and DksA discussed in the text are labeled.

**Supplemental Figure 1.**
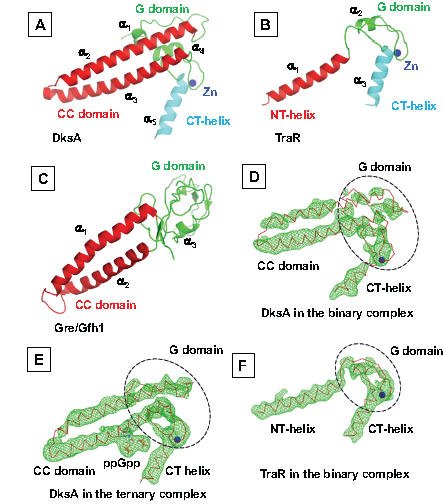
Structures of DksA, TraR and Gre/Gfh1 and *Fo-Fc* density maps showing DksA, DksA/ppGpp and TraR in their complexes with RNAP. (**A-C**) The crystal structures of apo-form DksA (PDB: 1TLJ, molecule A) (**A**), TraR in the RNAP–TraR complex (PDB: 5W1S) (**B**) and Gre/Gfh1 chimera in the RNAP–Gre/Gfh1 complex (PDB: 4WQT) (**C**). Structural parts of these factors and positions of α helixes are indicated. (**D-F**) *Fo-Fc* electron density maps (green mesh, σ=3) of the DksA in the binary (**D**) and ternary complexes (**E**) and the TraR in the RNAP–TraR complex (**F**). DksA and TraR are shown as red backbone ribbons, Zn atoms in their Zn binding site are blue sphere and ppGpp is depicted as a stick model. Three structural parts of DksA and TraR are indicated.

**Supplemental Figure 2.**
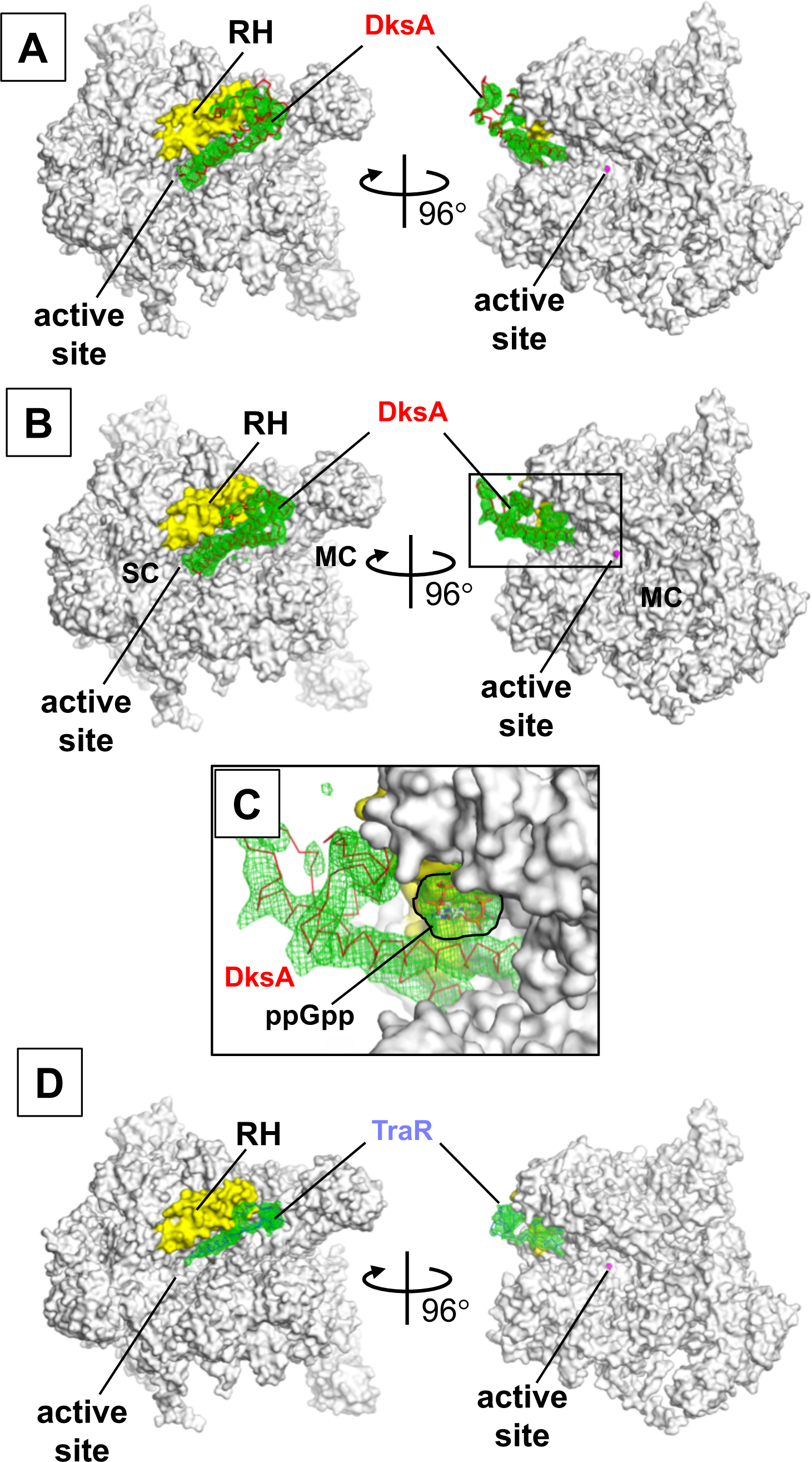
*Fo-Fc* density maps showing DksA, DksA/ppGpp and TraR in their complexes with RNAP. *E. coli* RNAP holoenzyme is depicted as molecular surface (white) with the active site Mg (magenta sphere) and β’rim helix (yellow) shown in colors. *Fo-Fc* electron density maps (σ=3) of RNAP—DksA (**A**) and RNAP—DksA/ppGpp (**B**) complexes phased with the holoenzyme are shown as green meshes. The DksA is shown as red backbone ribbon and ppGpp is depicted as stick model. The right and left panels show the main channel (MC) and the secondary channel (SC) of RNAP in their middles, respectively. (**C**) A magnified view of the boxed region in **B**, showing a *Fo-Fc* map corresponding to ppGpp (outlined by black). (**D**) *E. coli* RNAP holoenzyme is depicted as molecular surface (white) with the active site Mg (magenta sphere) and β’rim helix (yellow) shown in colors. *Fo-Fc* electron density maps (σ=3) of the RNAP–TraR complex phased with the holoenzyme are shown as green meshes. TraR is shown as a backbone ribbon.

**Supplemental Figure 3.**
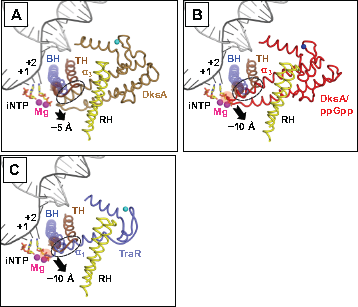
The acidic tips of DksA and TraR restrict trigger loop folding and the iNTP binding at +2 position. DksA and TraR are depicted as ribbon models in **(A)** the RNAP—DksA complex, **(B)** the RNAP—DksA/ppGpp complex and **(C)** the RNAP–TraR complex. The β’rim helix and bridge helix of RNAP are also depicted. The folded trigger loop, the trigger helix (TH), is modeled from the *T. thermophilus* transcription elongation complex (PDB: 2O5J), while the DNA and iNTPs are modeled using the *T. thermophilus* transcription initiation complex (PDB: 4Q4Z). Movements of the CC tip of DksA and the NT-helix of TraR required for folding the TH are indicated by arrows.

**Supplemental Figure 4.**
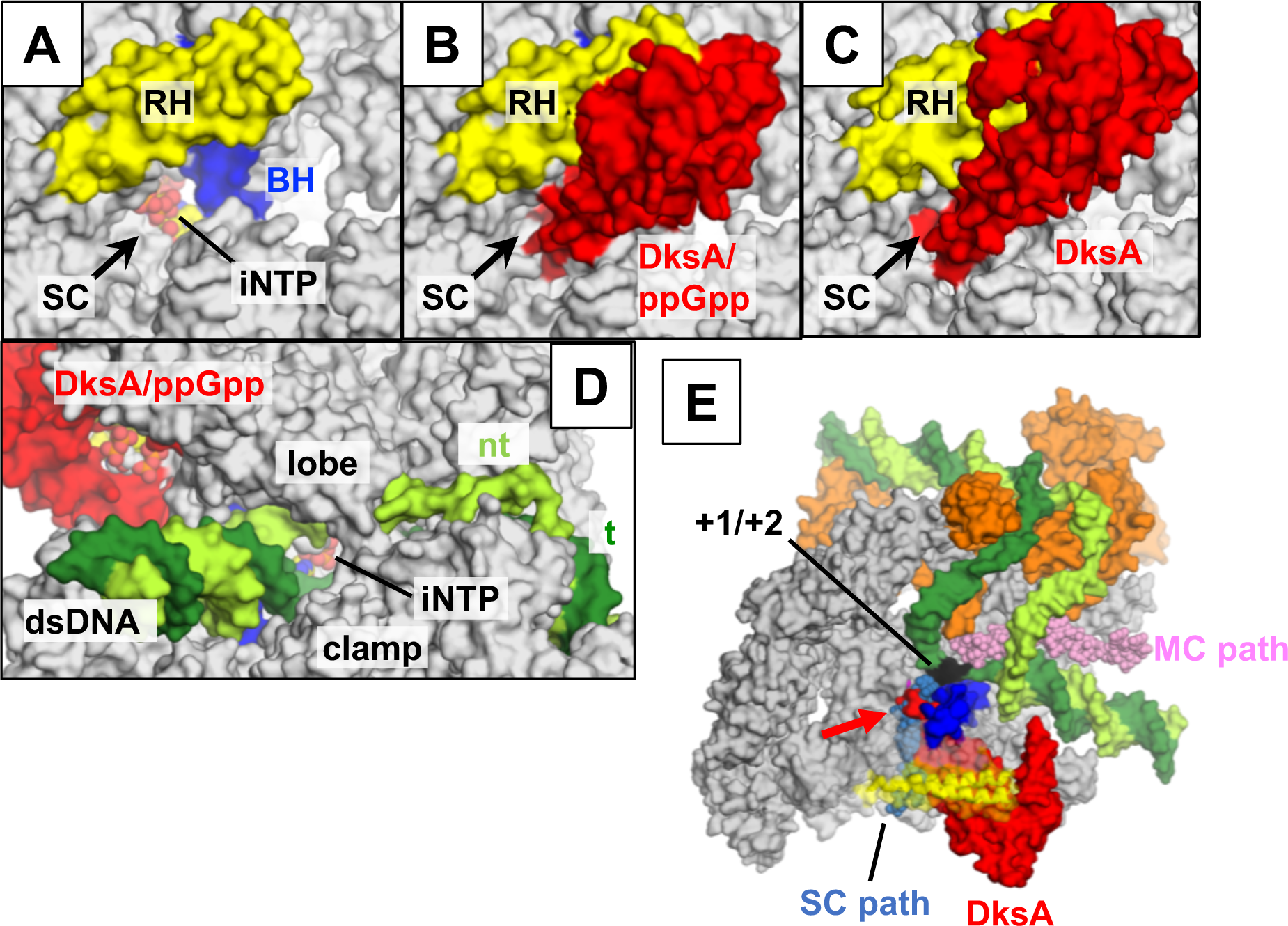
RNAP and DksA interaction and the NTP entry routes to the open promoter complex in the presence of DksA or TraR. **(A-D)** The iNTP (CPK representation) bound at the active site are viewed through the secondary channel **(A:** apo-form RNAP, **B:** RNAP—DksA/ppGpp, **C:** RNAP—DksA) and the main channel (D RPo with DksA/ppGpp). Molecular surfaces of RNAP (white), DksA (red) and DNA (template DNA: dark green, non-template DNA: light green) are depicted, and the β’rim helix (yellow) and bridge helix (blue) of RNAP are highlighted. (E) The main channel can be used for NTP entry of the RPo with DksA covering at the secondary channel. RNAP, DNA and DksA are depicted as surface, CPK and ribbon representations, β subunit is removed to see the main channel of RNAP. NTP molecules aligned along the NTP entry route from the main channel are shown in pink with CPK representation.

**Supplemental Figure 5.**
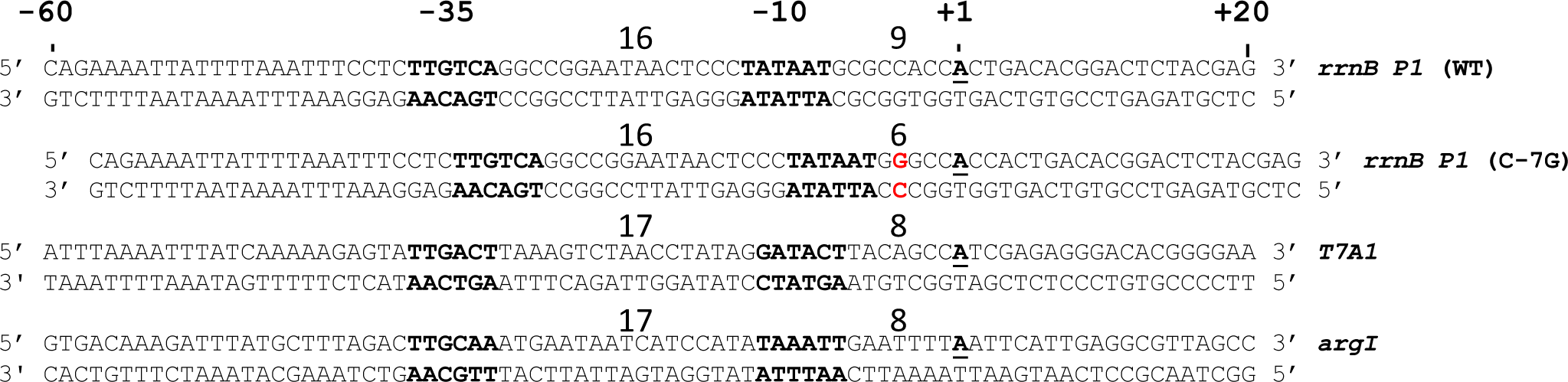
Promoter DNA sequences used for the measurement of the DksA binding to RNAP – promoter DNA complex **(Fig. 5).** Lengths between the -35 and -10 elements as well as distances between the downstream end of −10 element to the transcription start site of each promter are indicated.

**Supplemental Figure 6.**
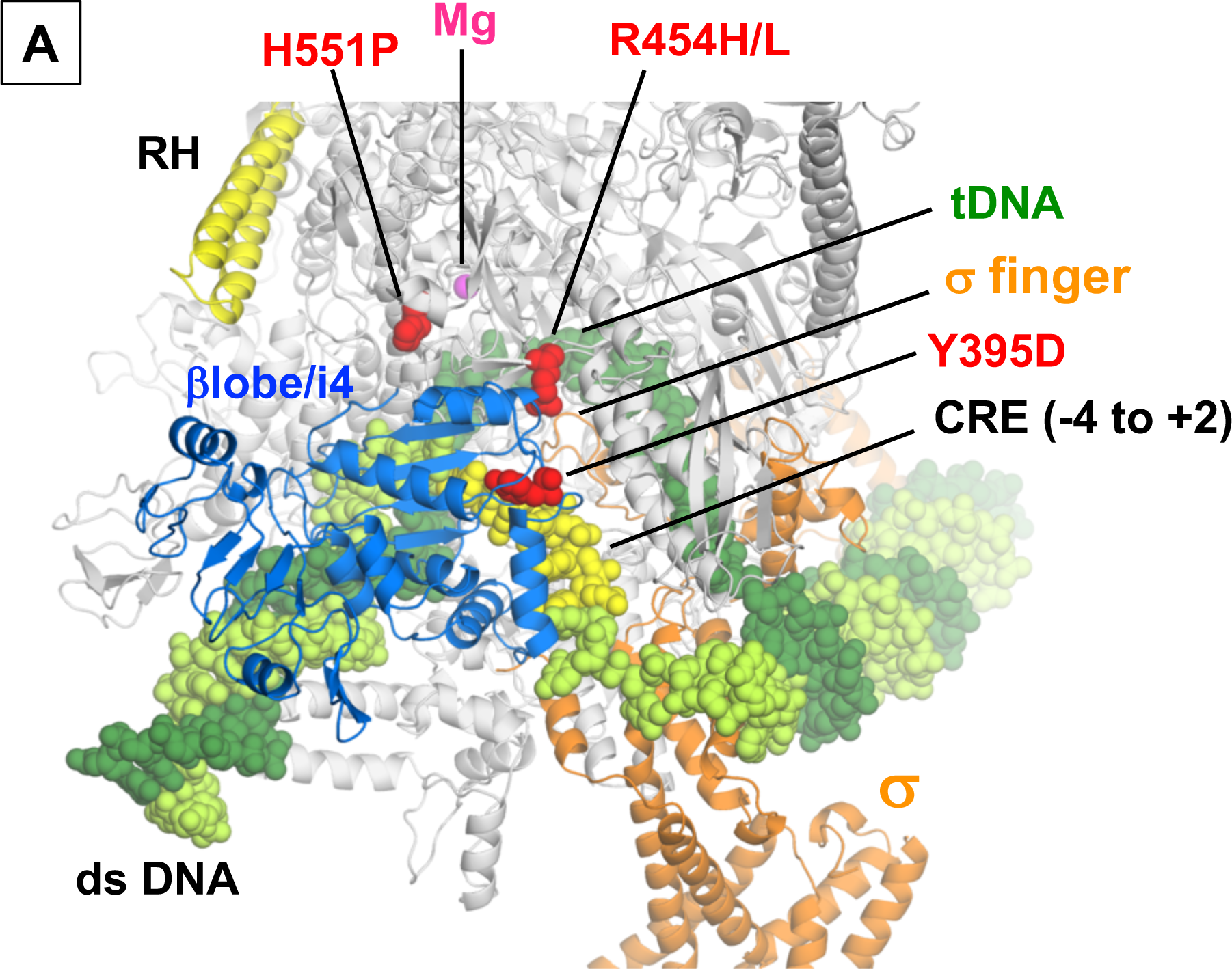
The RNAP open promoter complex model. **(A)** *E. coli* σ^70^ holoenzyme and open complex DNA model are depicted as ribbon and CPK representations, respectively. RNAP domains, σ subunit and core recognition element (CRE) of the non-template DNA are highlighted and labeled. *ΔdksA* suppressors (Rutherford et al., 2009) are shown as spheres and labeled.

**Supplemental Figure 7.**
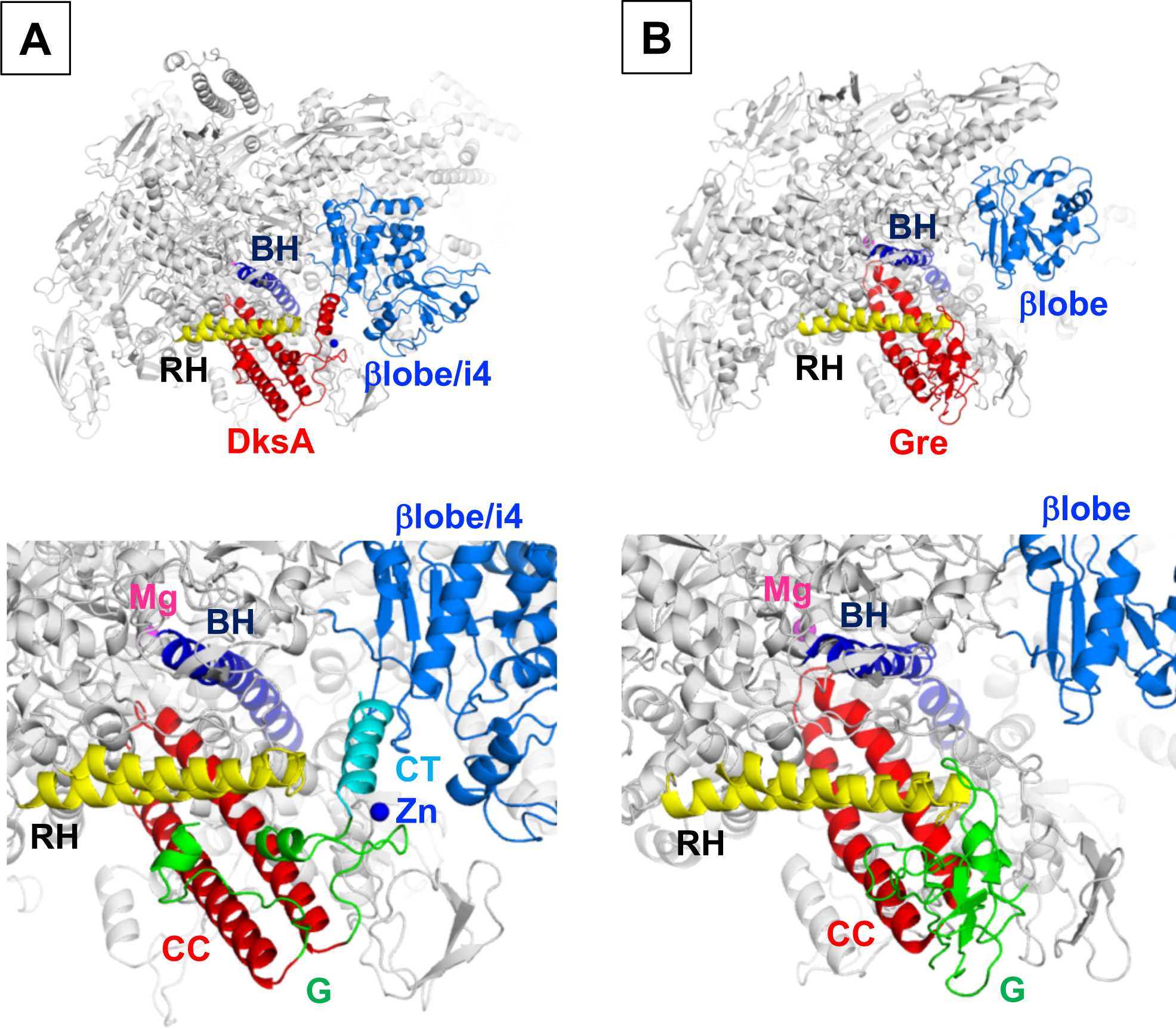
Comparison between the DksA and Gre modes of binding to RNAP. Structures of the *E. coli* RNAP and DksA complex **(A)** and the *T. thermophilus* RNAP and Gre/Gfh1 chimera protein complex structures **(B)** (PDB: 4WQT). RNAP, DksA and Gre/Gfh1 are depicted as ribbon models. DksA and Gre/Gfh1 binding regions are magnified in the panels below and structural parts of DksA and Gre/Gfh1 chimera are indicated. The orientations of panels are the same as in a right panel of **Fig. 1A.**

**Supplemental Figure 8.**
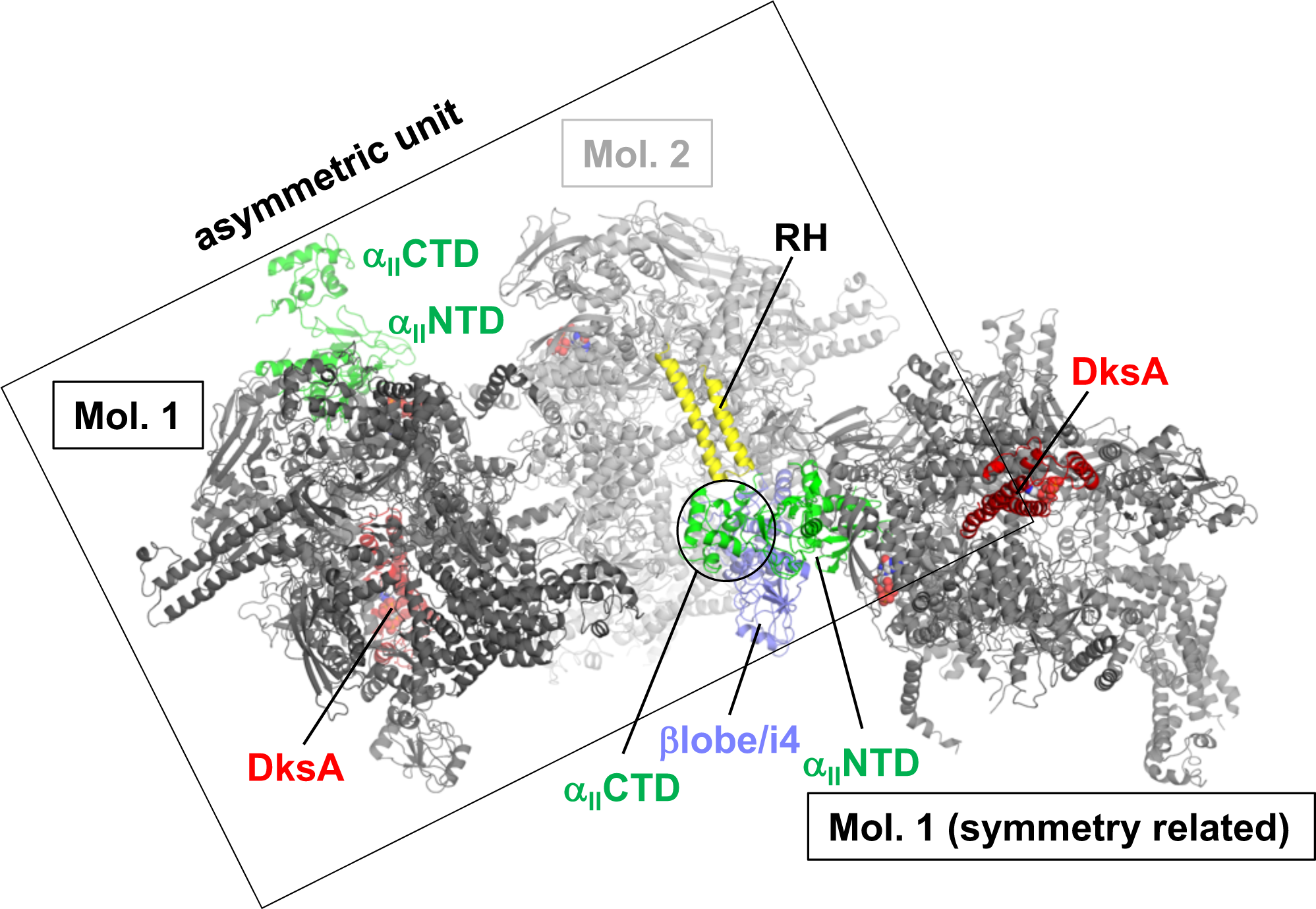
α_II_ CTD of the first RNAP molecule occupies the DksA binding site of the second RNAP molecule in crystal, which prevents the DksA binding to the second RNAP molecule.

